# Stretching of the retinal pigment epithelium contributes to zebrafish optic cup morphogenesis

**DOI:** 10.1101/2020.09.23.310631

**Authors:** Tania Moreno-Mármol, Mario Ledesma-Terrón, Noemí Tabanera, María Jesús Martin-Bermejo, Marcos J Cardozo, Florencia Cavodeassi, Paola Bovolenta

## Abstract

The vertebrate eye primordium consists of a pseudostratified neuroepithelium, the optic vesicle (OV), in which cells acquire neural retina or retinal pigment epithelium (RPE) fates. As these fates arise, the OV assumes a cup-shape, influenced by mechanical forces generated within the neural retina. Whether the RPE passively adapts to retinal changes or actively contributes to OV morphogenesis remains unexplored. Here, we generated a zebrafish Tg(E1-*bhlhe40*:GFP) line to track RPE morphogenesis and interrogate its participation in OV folding. We show that, in virtual absence of proliferation, RPE cells stretch into a squamous configuration, thereby matching the curvature of the underlying retina. Forced proliferation and localized interference with the RPE cytoskeleton disrupt its stretching and OV folding. Thus, extreme RPE flattening and accelerated differentiation are efficient solutions adopted by fast-developing species to enable timely optic cup formation. This mechanism differs in amniotes, in which proliferation largely drives RPE expansion with a much-reduced need of cell flattening.

## Introduction

The retinal pigment epithelium (RPE) is an essential component of the vertebrate eye, composed of a monolayer of pigment-enriched epithelial cells abutting the neural retina (NR) with a primary role in photoreception (1). Despite the acquisition of specialized epithelial properties, RPE cells have a neural origin and share progenitors with the NR. These progenitors are organized in a pseudostratified neuroepithelium, known as optic vesicle (OV) or eye primordium. In amniotes, the OVs appear as balloon structures positioned at the sides of the anterior neural tube (2). In zebrafish instead, these primordia are flat and form two bilayered structures with the outer and inner layers distally connected by a rim or hinge (3). Under the influence of inductive signals (4, 5), the two layers activate different genetic programs that specify the cells of the inner layer and ventral outer layer as NR and those of the dorsal outer layer as RPE (6–8). Whilst this specification occurs, the OV bends assuming a cup-like shape (9).

The discovery of the *ojoplano* medaka fish mutant in which the OV remains unfolded, was instrumental to propose that basal constriction of NR progenitors is at the basis of OV bending (10). This basal constriction is mediated by the redistribution of the actomyosin cytoskeleton (10–12), which also enables the apical relaxation of retinal cells (13), enhanced by focal adhesions of the apical surface with the extracellular matrix molecules (ECM) such as laminin (12). The importance of concomitant apical relaxation, especially of the cells positioned at the hinge, has been also supported in studies of mammalian retinal organoids (14, 15). Nevertheless and independently of their relative contribution, the acquisition of apical convexity and basal concavity in the NR epithelium are accepted drivers of the biomechanical forces that induce OV folding (15). In zebrafish, this mechanism is reinforced by rim involution or epithelial flow, a process whereby progenitors at the hinge emit dynamic lamellipodia at the basal side and actively translocate from the ventral outer layer of the OV into the inner/retinal layer (3,13,16–19). Periocular neural crest cells appear to facilitate this flow, in part by the deposition of the ECM (20) to which the lamellipodia attach (13,17,18). The result of this flow is an imbalanced cell number between the two layers, which should favor NR bending (13,17,18) Whether this flow may also contribute to the concomitant cell shape modifications that the remaining outer layer cells undergo as they become specified into RPE, or conversely whether RPE specification favors the flow (17) remain open questions.

Indeed as the OV folds, the pseudostratified neuroepithelial cells of the OV dorsal outer layer progressively align their nuclei becoming a cuboidal monolayer in amniotes species (2, 21). In zebrafish, cuboidal cells further differentiate to a flat/squamous epithelium (16, 18) that spreads to cover the whole apical surface of the NR (16, 22). In mice, failure of RPE specification, as observed after genetic inactivation of key specifier genes (i.e. *Otx1/Otx2, Mitf, Yap/Taz*), enables RPE progenitors to acquire a NR fate (23–25). The resulting optic cups (OCs) present evident folding defects (23), raising the possibility that specific RPE features are needed for OC formation. In line with this idea, a differential stiffness of the RPE versus the NR layer has been proposed to drive the self-organization of mammalian organoids into an OC (14,15,26). Furthermore, generation of proper RPE cell numbers seems a requirement for correct OC folding in mice (27). However, studies addressing the specific contribution of the RPE to OV folding are currently lacking.

Here we report the generation of a Tg(E1-*bhlhe40*:GFP) zebrafish transgenic line with which we followed the beginning of RPE morphogenesis under both normal and interfered conditions. We show that, whereas in amniotes, including humans, the developing RPE undergo proliferation to increase its surface with a less evident cell flattening, zebrafish RPE cells rapidly cease proliferation and expand their surface by reducing their length along the apico-basal axis and extending in the medio-lateral direction with a cell-autonomous process that depends on cytoskeletal reorganization. Localized interference with either the retinal or the RPE actomyosin and microtubule cytoskeleton shows that RPE flattening actively contributes to OV folding. This represents an efficient solution to match the increased apical surface of the NR layer in a fast-developing vertebrate species such as zebrafish.

## Results

### Generation of a specific reporter line to study zebrafish RPE development

Detailed analysis of zebrafish RPE morphogenesis has been hampered by the lack of a suitable transgenic line, in which RPE cells could be followed from their initial commitment. The E40 (*bhlhe40*) gene, a basic helix-loop-helix family member, encodes a light and hypoxia-induced transcription factor (also known as *Dec1, Stra13, Sharp2* or *Bhlhb2*) involved in cell proliferation and differentiation as well as in the control of circadian rhythms (28). In neurulating zebrafish embryos, its expression is limited to cells of the prospective RPE (Fig. 1A) (22, 29), representing a potentially suitable tissue marker.

**Figure 1.**
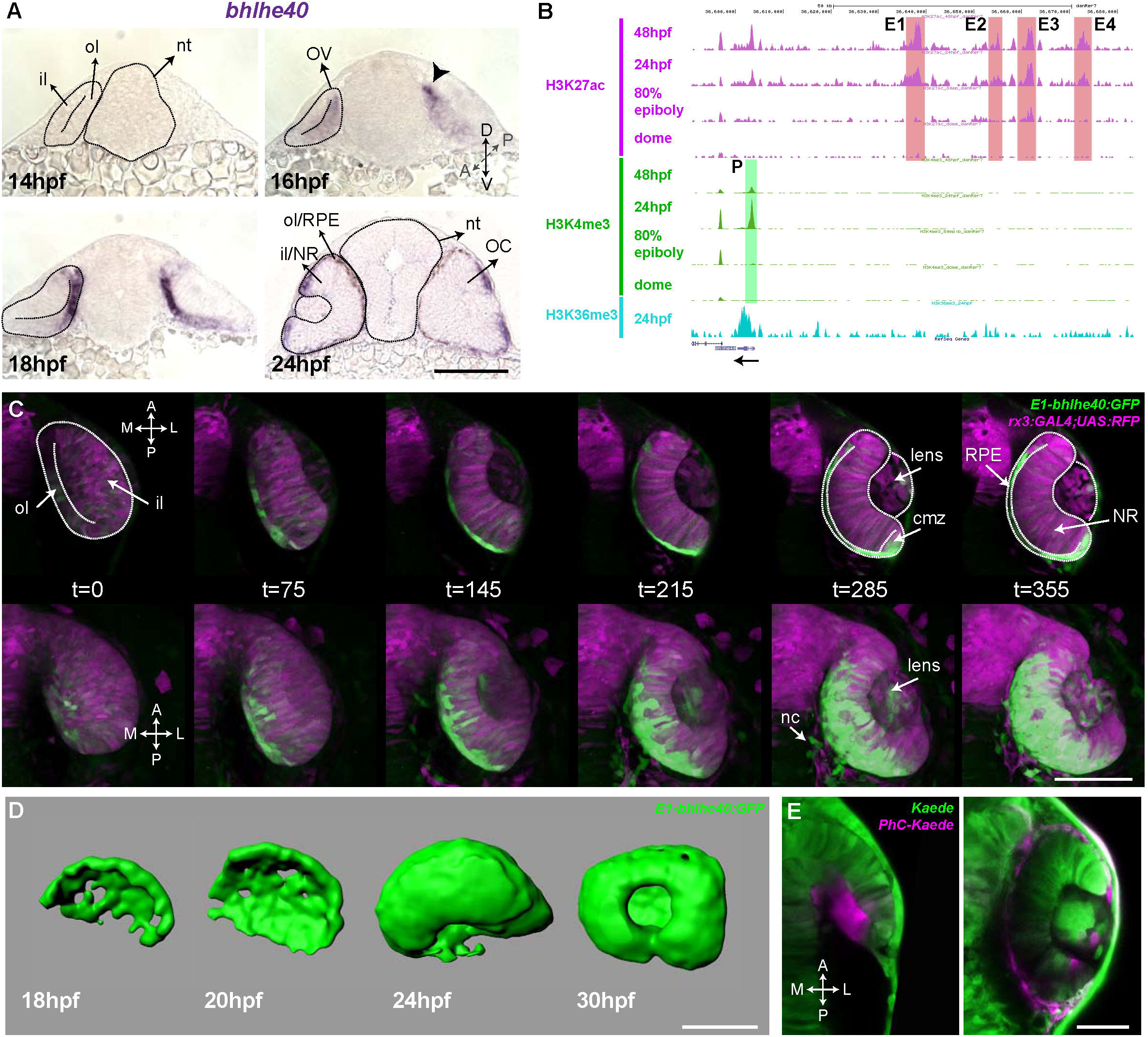
The Tg(E1-*bhlhe40*:GFP) line is a suitable tool to study early RPE generation. **A**) Frontal cryo-sections of 14-24 hpf wt embryos hybridized *in toto* with a *bhlhe40* specific probe. mRNA is first detected in the dorsal most region of the OV outer layer (arrowhead) and then expands ventrally. **B**) UCSC Genome Browser view of H3K27ac (purple, potential active enhancers), H3K4me3 (green, potentially active promoters) and H3K36me3 (light blue, transcriptionally active regions) tracks obtained for four zebrafish developmental stages: dome, 80% epiboly, 24 hpf, 48 hpf related to the upstream *bhlhe40* genomic locus (50kb). The black arrow at the bottom indicates *bhlhe40* position and direction. The promoter (P) and the four selected enhancers (E1 to E4) are highlighted with a color-coded box. **C)** Timeframes from *in vivo* time-lapse recording of a Tg(E1-*bhlhe40*:GFP;*rx3*:GAL4;UAS;RFP) embryo between 14-24 hpf. Time is indicated in min. Note that the GFP reporter signal matches the *bhlhe40* mRNA distribution in A. **D)** 3D reconstruction of the prospective RPE from Tg(E1-*bhlhe40*:GFP) embryos at the stages indicated in the panel. **E)** Dorsal view of a wt embryo injected with Kaede mRNA (green) at 12 hpf. A group of cells in the dorsal region of the outer layer was photoconverted (magenta, panel on the left) and the embryo visualized at 30 hpf (right panel). Magenta labelled cells cover the entire RPE region. Black and white dashed lines delineate the OV, neural tube and virtual lumen in **A, C.** Abbreviations: A, anterior; cmz, ciliary margin zone; il, inner layer; l, lateral; m, medial; NR, NR; OC, OC; ol, outer layer; OV, optic vesicle; P, posterior; RPE, retinal pigment epithelium. Scale bars: 100 μm (A-D); 50 μm, E.

We used predictive enhancer and promoter epigenetic marks at different zebrafish developmental stages (30) to scan the *bhlhe40* locus for the presence of conserved and active regulatory regions. The promoter and four potential enhancers (E1-4; Fig. 1B) appeared to be active between 80% epiboly and 24 hpf, encompassing the early stages of zebrafish eye development (30). These enhancers were selected, amplified and tested using the ZED vector (31) as potential drivers of gene expression in the prospective RPE. The resulting F0 embryos were raised to adulthood and screened. Only the E1 enhancer drove specific and restricted GFP reporter expression into the prospective RPE. The corresponding fishes were further crossed to establish the stable transgenic line Tg(E1-*bhlhe40*:GFP) used in this study.

Time-lapse studies of the Tg(E1-*bhlhe40*:GFP) progeny confirmed that the transgenic line faithfully recapitulated the *bhlhe40* mRNA expression profile detected with *in situ* hybridization (Fig 1A,C). GFP reporter expression appeared in a discrete group of neuroepithelial cells in the dorso-medial region of the OV (16-17 hpf) and expanded both posteriorly and ventrally (Fig 1C; Movie S1 and S2), so that, by 24 hpf, GFP positive cells appeared to wrap around the entire inner NR layer. 3D reconstructions of selected embryos further confirmed the fast (about seven hours) expansion of the GFP-positive domain forming an outer shell for the eye (Fig. 1D). During this process no GFP expression was observed in regions other than the RPE. However, after the formation of the OC, reporter expression appeared also in the ciliary marginal zone (CMZ), the pineal gland and few neural crest cells surrounding the eye (Fig. 1C; Movie S1-3). These additional domains of expression coincided with the reported *bhlhe40* mRNA distribution (29) and represented no obstacle for using the transgenic line as a tool to follow the early phases of RPE generation. Indeed, very early activation represents an important advantage of the Tg(E1-*bhlhe40*:GFP) line over other presently available transgenic lines that allow visualizing de RPE (32, 33).

The suitability of the Tg(E1-*bhlhe40*:GFP) line for the identification of the very first RPE cells is supported by the onset of the reporter expression in the dorso-medial OV region, coinciding with previous fate map predictions (16, 18). To further verify this notion, we took advantage of the characteristic of the fluorescent Kaede protein (34) that switches from green to red emission upon UV illumination. Embryos were injected with Kaede mRNA and neuroepithelial cells located at the most dorso-medial region of the OV were UV illuminated at the 15 hpf stage to ensure that no differentiation had yet occurred (Fig. 1E). Embryos were let develop until 30 hpf. Photoconverted cells were found throughout the thin outer layer of the OC (Fig. 1E), confirming that the entire RPE derives from the dorso-medial OV region.

### Neuroepithelial cell flattening drives RPE expansion at OV stages

Tg(E1-*bhlhe40*:GFP) embryos were thereafter used to dissect the extensive changes in cell shape that are associated with the acquisition of RPE identity (16, 22). At OV stage all retinal progenitors present a columnar-like morphology characteristic of embryonic neuroepithelia (Fig. 2A,A’). As soon as RPE progenitors begin to express the transgenic GFP reporter, their apico-basal length rapidly and progressively reduces (Fig. 2A-C’), so that the cells first assume a cuboidal shape (Fig. 2B,B’) and then become flat, forming a squamous epithelial monolayer overlaying the apical surface of the NR (Fig. 2C,C’). At 30 hpf, RPE cells presented a polygonal, frequently hexagonal, morphology (Fig 2D,D’), with an apical surface area that, on average became about eightfold larger than that observed in progenitor cells (Fig. 2F; RPE ā: 354.8 ± 100.3 μm^2^ vs PN ā: 43.7 ± 7.8 μm^2^). In contrast, the abutting apical surface of NR cells slightly shrank as compared to that of progenitor cells (Fig. 2E,E’,F; NR ā: 22.5 ± 2.9 μm^2^ vs PN ā: 43.7 ± 7.8 μm^2^) while maintaining a constant apico-basal length. The latter observation agrees with previous reports showing that the cone-like morphology of NR progenitors represents only a slight modification of the progenitor columnar shape (11, 13).

**Figure 2.**
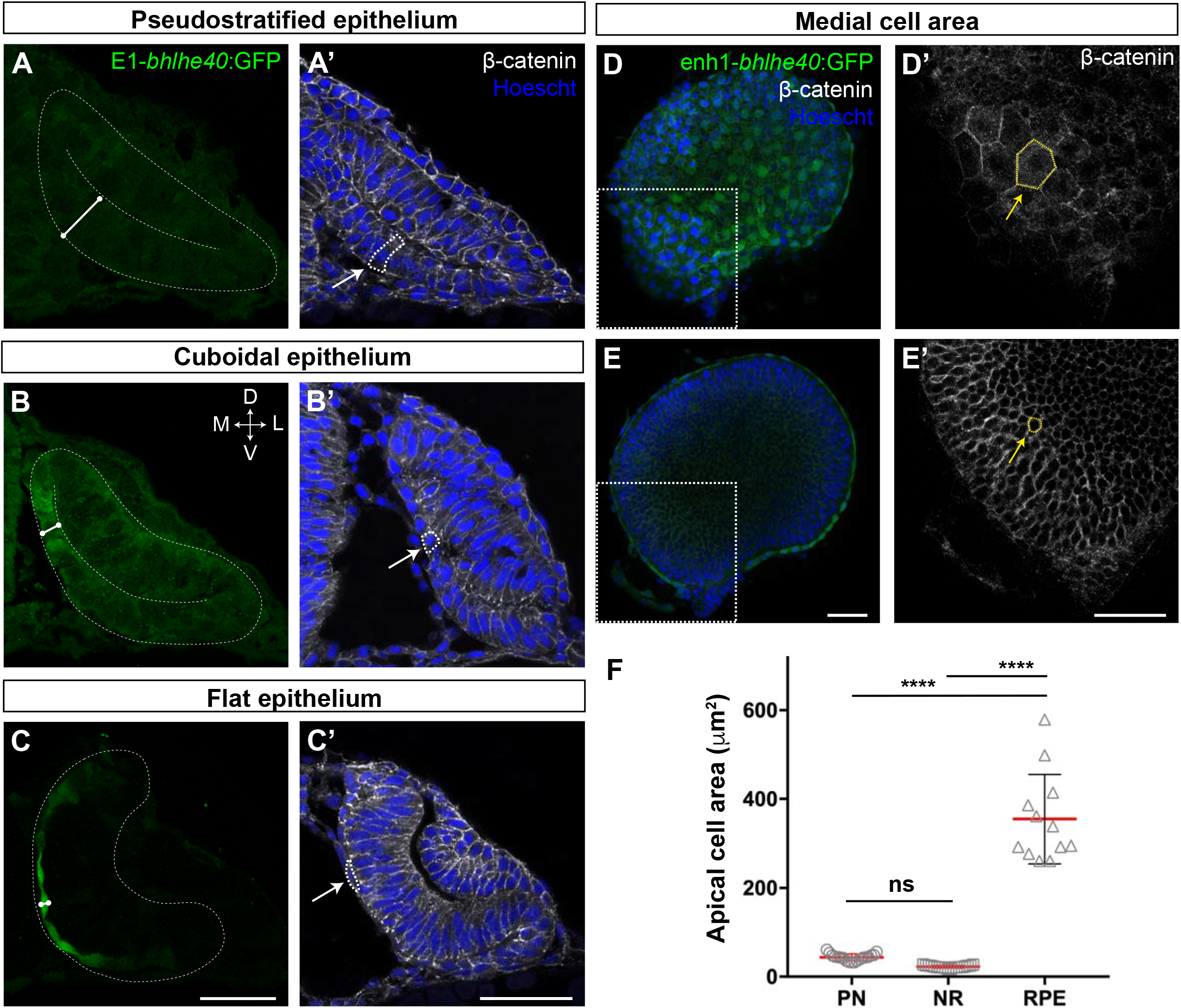
The RPE converts from a pseudostratified to a squamous epithelium during OV folding by increasing individual cell surface. **A-C’)** Confocal images of frontal cryosections of Tg(E1-*bhlhe40*:GFP) embryos immunostained for GFP (green) and β-catenin (white) and counterstained with Hoechst (blue). Note that the RPE rapidly decreases its thickness (white straight line in **A-C**) and cells change from columnar (arrow in **A’**) to cuboidal (arrow in **B’**) and then flat shape (arrow in **C’**). White dashed lines delineate eye contour and virtual lumen in A-C. **D-E’**) Confocal images of the posterior RPE (D, D’) and NR (E, E’) regions of an eye cup dissected from 30 hpf Tg(E1-*bhlhe40*:GFP) embryos immunostained for GFP (green) and β-catenin (white) and counterstained with Hoechst (blue). Images in D’, E’ are high power views of the areas boxed in white box in D, E. Note the hexagonal morphology (yellow arrow in D’) of RPE cells (average area 354.8 ± 100.3 μm^2^) in contrast to the small and roundish cross-section of retinal progenitors (average area 22.5 ± 2.9 μm^2^; yellow arrow in E’). **F**) The graph represents the average area of individual OV progenitors and NR and RPE cells. The average area is calculated using cells from 5 different embryos. Data represent mean ± SD, **** p<0.0001. ns non-significant. Scale bar: 50 μm.

To obtain a quantitative analysis of the dynamic changes that RPE tissue, as whole, underwent during OV folding, we performed a morphometric characterization of the images from movies S1-3. To this end, the fluorescent information from the Tg(E1-*bhlhe40*:GFP) reporter was discretized into seven different segments that were individually analyzed along the recording time (Fig. S1; material and methods). The combined quantification of the different segments (Fig. S1, Fig. 3A) showed that, between stages 17 and 21 hpf, the overall thickness of the RPE tissue underwent, on average, a flattening of more than threefold (from a mean of about 24 to 8 μm; Fig. 3B). Flattening occurred with a central to peripheral direction, so that RPE cells closer to the hinges were the last ones to flatten (Fig. 2C,C’). In parallel, the overall RPE surface underwent a ~ two-fold expansion between 17 and 22 hpf (from approx. 1.1 to 2.2 x 10^3^ μm^2^; Fig. 3C; Movie S1-2), reflecting the large apical area increase observed in each individual cell at later stages (Fig. 2F). In line with the idea that cell flattening is *per se* sufficient to account for whole tissue enlargement, the RPE volume only slightly changed between 17 and 20 hpf with a slope increase of 0.47 x 10^3^ μm^3^/h (Fig 3D).

**Figure 3.**
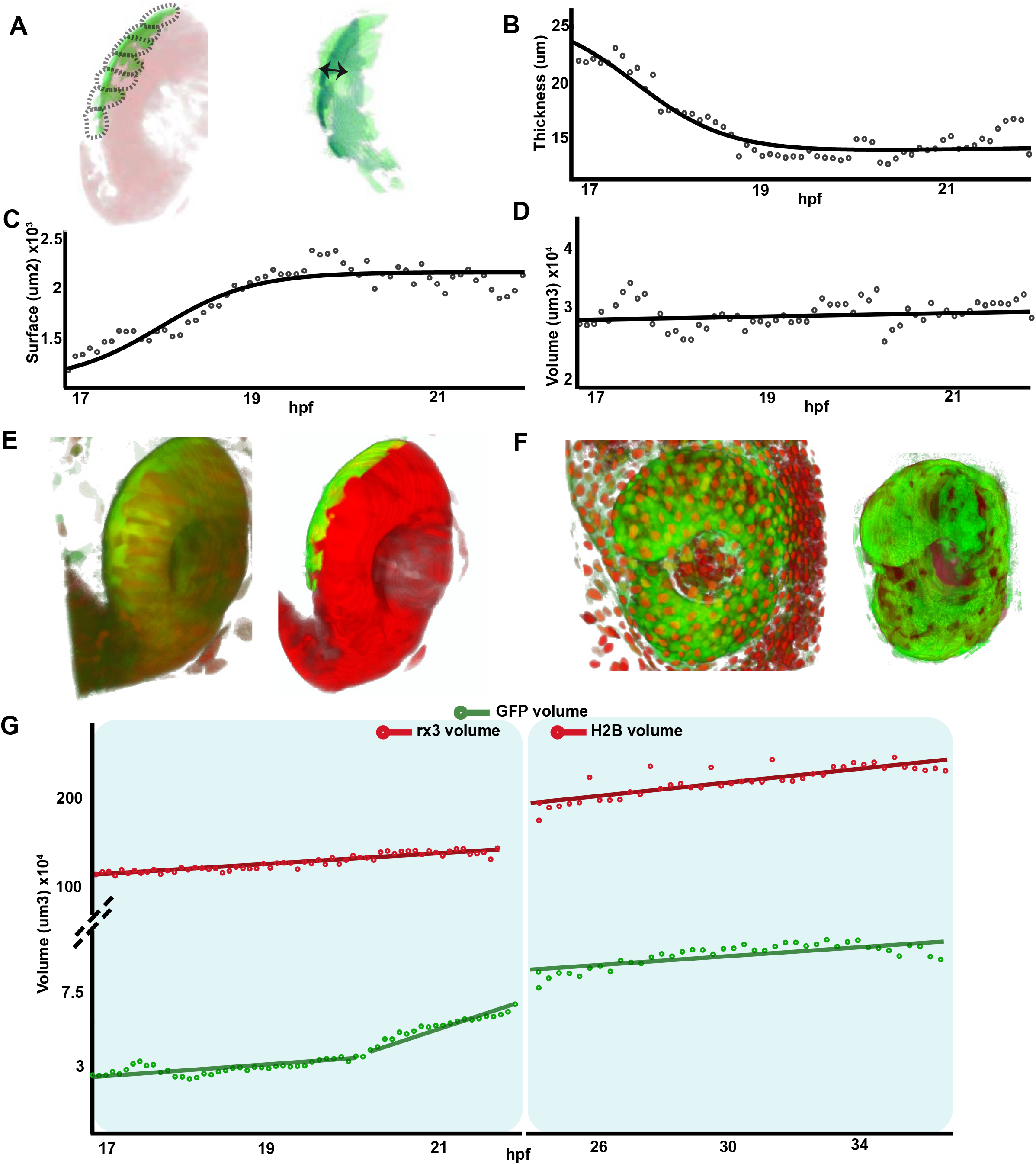
RPE volume is conserved during initial tissue morphogenesis. **A)** Image on the left represents the reconstruction of a single frame from movie S2 showing the OV/OC in red and the RPE in green. The segments in which the RPE was discretized are depicted with black dashed lines. The image on the right shows the RPE reconstruction obtained after filtering. Double arrow points to RPE thickness. **B-D**) The graphs show how the RPE thickness (B, calculated as volume/surface), surface (C) and volume (D) change as a function of the developmental stage. **E**) 3D reconstructions of raw (left) and processed (right) versions of a frame from the Movie S1-2. **F**) 3D reconstructions of raw (left) and processed (right) versions of a frame from the Movie S3. **G**) Quantification of RPE and eye volume based on Movie S1-2 (rx3 volume quantification) and the Movie S3 (H2B volume quantification) along developmental stages.

To provide further support to this idea we analyzed the RPE volume variation in comparison with the growth of the entire OC. To this end, we analyzed RPE and OC volume in two time windows: from 17 to 22 hpf (Movie S1-3) and from 24 to 37 hpf (Movie S4), using GFP (RPE) and RFP (eye) reporter signals from the double Tg(E1-*bhlhe40*:GFP; *rx3*:GAL4;UAS;RFP) line or from the Tg(E1-*bhlhe40*:GFP) line injected with the *pCS2:H2B-RFP* mRNA (Fig. 3E,F). Signal quantification showed that the eye underwent a marked and linear volume increase (slope: 5.54 x 10^4^ μm^3^/h from 17 to 22 hpf and 3.6 x 10^4^ μm^3^/h from 24 to 37 hpf) as compared to that of the RPE (Fig. 3D,G). Nevertheless, RPE reporter volume suddenly expanded between 20 and 22 hpf (slope: 1.25 x 10^4^ μm^3^/h; Fig. 3G) to then slow back between 24 to 37 hpf (slope: 1.2 x 10^3^ μm^3^/h; Fig. 3G). Closer examination of the timelapse movies showed that the sudden peak of GFP-positive volume expansion corresponded to reporter activation in the posterior and, to a lesser extent, in the anterior CMZ (Movie S1-2). Confirming this association, only the tissue segments very close to the posterior CMZ had a volume larger than that of the RPE at 17-20 hpf (Fig. S1), whereas RPE segments located in the most central regions presented a volume undistinguishable from that detected at previous stages. In sum, a comparison of the dynamics slopes from central RPE and OV regions suggests that the volume of the RPE grows at very low pace (0.47 x 10^3^ μm^3^/h) – despite the rather drastic morphological changes of its cells –, whereas the whole OV expands at a pace ~ twenty-five times faster (1.25 x 10^4^ μm^3^/h; Fig. 3D)

Taken all together, this morphometric analysis indicates that the expansion of the RPE in zebrafish occurs by the recruitment of a limited number of cells that undergo profound cell shape changes: from a neuroepithelial to squamous morphology.

### RPE flattening is a cell autonomous process required for proper OV folding

Both external interactions and intracellular processes determine the shape of a cell and define its mechanical properties (35). Thus, in principle, RPE flattening might occur as a “passive” process, triggered by the forces that the NR and hinge cells exert on the RPE (2, 17). Alternatively, it might depend on cell autonomous cytoskeletal rearrangements, involving, for example, myosin II activity, which controls the acquisition of a flat epithelial morphology in other contexts (36, 37). Discriminating between these two possibilities has been technically difficult. Experiments directed to assess the mechanisms of OV folding have used whole embryo bathing in drugs such as blebbistatin (11, 13), a specific myosin II inhibitor (38). Such an approach hampers the assessment of the potential influence of NR over RPE morphogenesis (and vice-versa) as well as the relative contribution of the two tissues to OV folding. We sought to overcome this limitation by spatially localized interference with the cytoskeletal organization of either the RPE or NR and by recording the cell autonomous and non-autonomous consequences. Nevertheless, to begin with, we reproduced the whole embryo bathing approach used by others (11, 13), focusing on the yet unreported effect that blebbistatin had on the RPE.

Tg(E1-*bhlhe40*:GFP) embryos were bathed either in blebbistatin or its diluent (DMSO) at 17 hpf (the onset of RPE specification; Fig 4A-C) and then let develop up to 19.5 hpf, when embryos were analyzed. DMSO-treated (control) embryos developed normally forming an OC surrounded by a squamous RPE (Fig. 4B). In blebbistatin-treated embryos, NR cells did not undergo basal constriction and the OV remained unfolded (Fig. 4C), as previously described (11, 13). Notably, in almost all the embryos analyzed (n=44/49) RPE cells did not flatten but remained cuboidal in shape (Fig. 4C). A similar phenotype was observed after treatment with para-nitro-blebbistatin, a non-cytotoxic and photostable version of blebbistatin (Fig. 4D), supporting that lack of OV folding is associated with alterations in both retina and RPE. To uncouple the two events we turned to the photoactivable compound azido-blebbistatin (Ableb), which binds covalently to myosin II upon two-photon irradiation, thus permanently interfering with myosin II activity in a spatially restricted manner as already proven (39, 40). Tg(E1-*bhlhe40*:GFP) 17 hpf embryos were bathed in Ableb or in DMSO and irradiated in a small region of either the dorsal outer layer (RPE) or the inner retinal layer (retina) of the OV (see methods). Embryos were then let develop until 24 hpf. Ableb photoactivation in the prospective RPE cells reproduced, although slightly less efficiently (n=30/44 embryos), the phenotype observed upon whole embryo bathing in blebbistatin, in which RPE cells acquired a cuboidal morphology (Fig. 4C,D,G,G’). No detectable alterations were found in the OV of irradiated/DMSO treated embryos or in the contralateral non-irradiated OV of embryos incubated in Ableb regardless of the irradiated region (Fig. 4E-F’, H-I’). Cell shape quantifications showed a significantly longer apico-basal axis (Fig. 4E’-G’) in irradiated Ableb RPE cells, normalized to that of control (DMSO and Ableb treated nonirradiated) OVs (Fig. 4K; Mann-Whitney U test, z=-5.088, p<0.001, control mean length 15.96 vs Ableb-treated 38.03). Failure of cell flattening in the irradiated region of the RPE was consistently associated with a significant reduction of OV folding (Fig. 4E-G), as assessed by measuring the invagination angle (13), which was normalized to that of control embryos (Fig 4L; Mann-Whitney U test: z=-2.704, p<0.01, mean rank for control 21.60 vs ABleb-treated 33.33). Photo-activation of Ableb in similar areas of the prospective NR basal region resulted in an elongated NR and a significantly impaired OV folding (Fig. 4J), as determined by the invagination angles normalized to those of control OV (Fig. 4N; Mann-Whitney U test: z=-3.035, p<0.01, mean rank control 10.29 vs Ableb 20.06). Notably, disruption of NR morphogenesis had no consequences on RPE development in all the analyzed embryos (n=16/22): cells underwent normal flattening with apico-basal lengths comparable to those of controls (Fig. 4H’-J’, M; Mann-Whitney U test: z=0.582, p>0.05, mean rank control 14.50 vs Ableb 16.38). This strongly supports that RPE flattening is not secondary to NR folding but rather a cell autonomous event. Notably, blebbistatin or Ableb treatments did not compromise the expression of the Tg(E1-*bhlhe40*:GFP) transgene in any experimental condition, indicating that cellular tension and morphology did not affect RPE specification.

**Figure 4.**
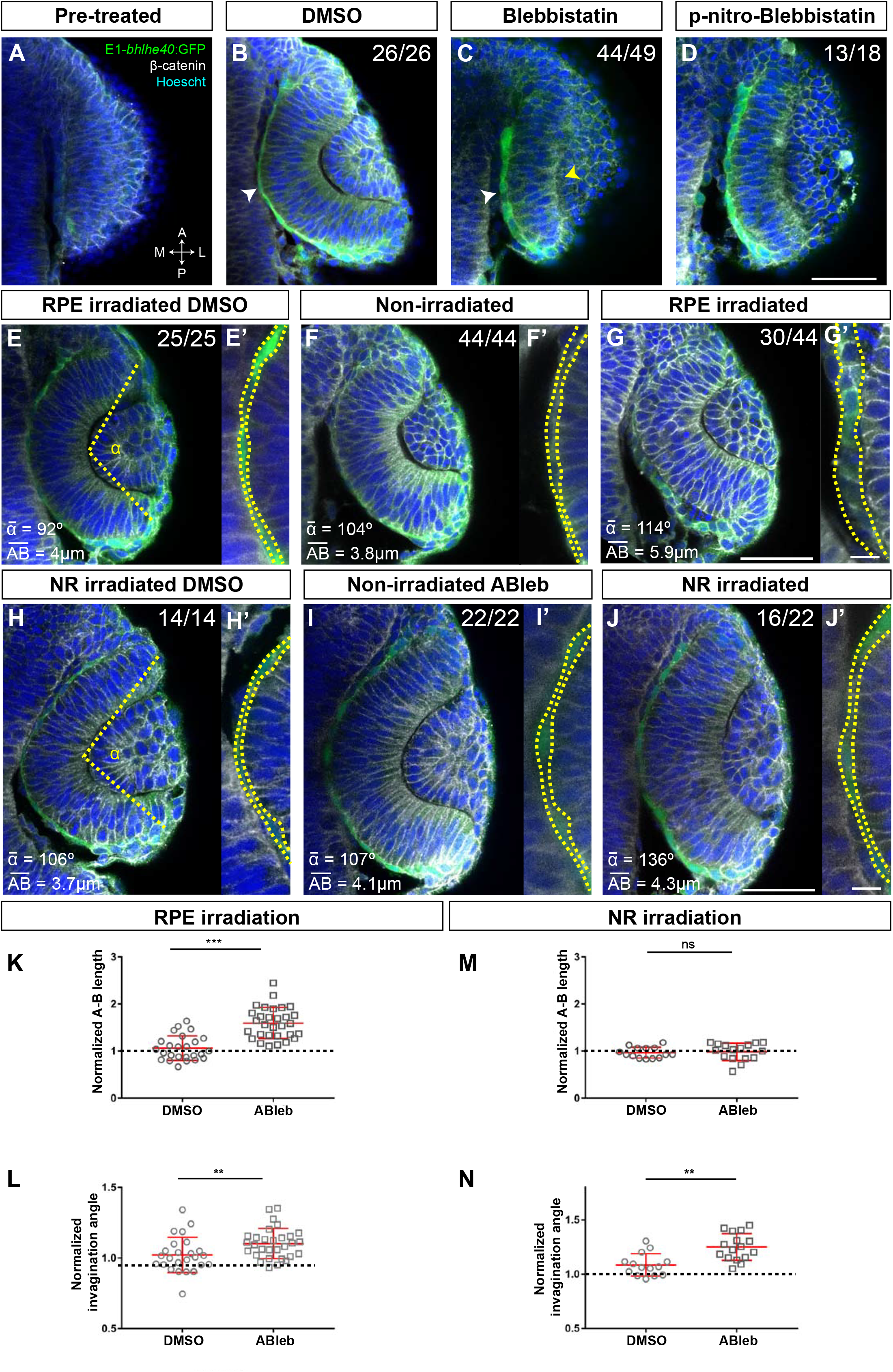
RPE flattening is a myosin-dependent cell-autonomous process required for proper OV folding. **A-J)** Confocal images of dorsally viewed Tg(E1-*bhlhe40*:GFP) embryos before (17 hpf; **A**) and 2.5 hr after incubation (19.5 hpf) with either DMSO (B, E, H), Blebbistatin (**C**), p-nitro-Blebbistatin (**D**) or ABleb (F, G, I, J) with (G, J) or without irradiation (F, I) in the prospective RPE (F-G) or NR (I-J). Images in E’, F’, G’ H’, I’ and J’ are high power views of RPE morphology. Embryos were immunostained for GFP (green), β-catenin (white) and counterstained with Hoechst (blue). Note that the OC forms and the RPE flattens (white arrowhead in B) normally in all DMSO treated embryos (B, E, E,’ H, H’) or in embryos incubated in Ableb without irradiation (F, F’, I, I’). In contrast, the RPE remains cuboidal (white arrowhead in C) and NR cells seem not undergo basal constriction (yellow arrowhead in C) in the presence of myosin inhibitors (C, D). Photoactivation of Ableb in the RPE prevents cell flattening (compare E’,F’ with G’) and impairs OV folding (G). When Ableb is photoactivated in the NR, folding of the OV is also impaired (J) but RPE cells undergo flattening (compare H’,I’ with J’). The number of embryos analyzed and showing the illustrated phenotype is indicated on the top right corner of each panel and the average invagination angle and mean A-B on the left bottom corner. The yellow dashed line in **E, H** indicate how the invagination angle (α) was determined. **(K, M)** Normalized RPE height in DMSO and ABleb treated embryos, irradiated either in the RPE (K) or in the NR (M). **(L, N)** Normalized invagination angle in DMSO and ABleb treated embryos irradiated either in the RPE (L) or in the NR (N). Data represent mean ± SD; ** p<0.01 and *** p<0.001. ns nonsignificant. Scale bars: 50 μm in A-J and 25 μm in E’-J.’

Microtubule dynamics has an important role in determining the cell shape (41). For example, reorientation of the microtubule cytoskeleton from the apico-basal to the medio-lateral cell axis together with actin filaments redistribution seems to drive the conversion of the *Drosophila* amnioserosa cells from a columnar to squamous epithelium (42). To determine if a similar reorientation occurs in the RPE we used time-lapse analysis of Tg(E1-*bhlhe40*:GFP) embryos injected with the mRNA of EB3:GFP, a protein that binds to the plus end of growing microtubules (43). In neuroepithelial RPE progenitors, microtubules grew in the apico-basal direction, whereas growth turned to the medio-lateral plane, as the RPE cells became squamous (Fig. S2; Movie S5-7). To determine if this reorientation is important for cell flattening, we bathed Tg(E1-*bhlhe40*:GFP) embryos in nocodazole, a drug that interferes with microtubule polymerization, or its vehicle (DMSO) at either 16 or 17 hpf and then analyzed them at 18.5 or 19.5 hpf, respectively. The eye of DMSO treated embryos developed normally (Fig. 5A, B), whereas in the presence of nocodazole RPE cells retained a columnar-like morphology with a stronger phenotype in embryos exposed to the drug at an earlier stage (Fig. 5C). Nocodazole treatment did not prevent the activation of the GFP reporter expression (Fig. 5C) or the acquisition/distribution of expected specification (*otx1* and *mitf*) and apico-basal polarity (zo-1 and laminin) markers (Fig S3A-H). Notably, although the NR layer appeared to bend inward, the RPE layer remained unfolded (Fig. 5F) and outer layer cells accumulated at the hinge, suggesting a defect in rim involution.

**Figure 5.**
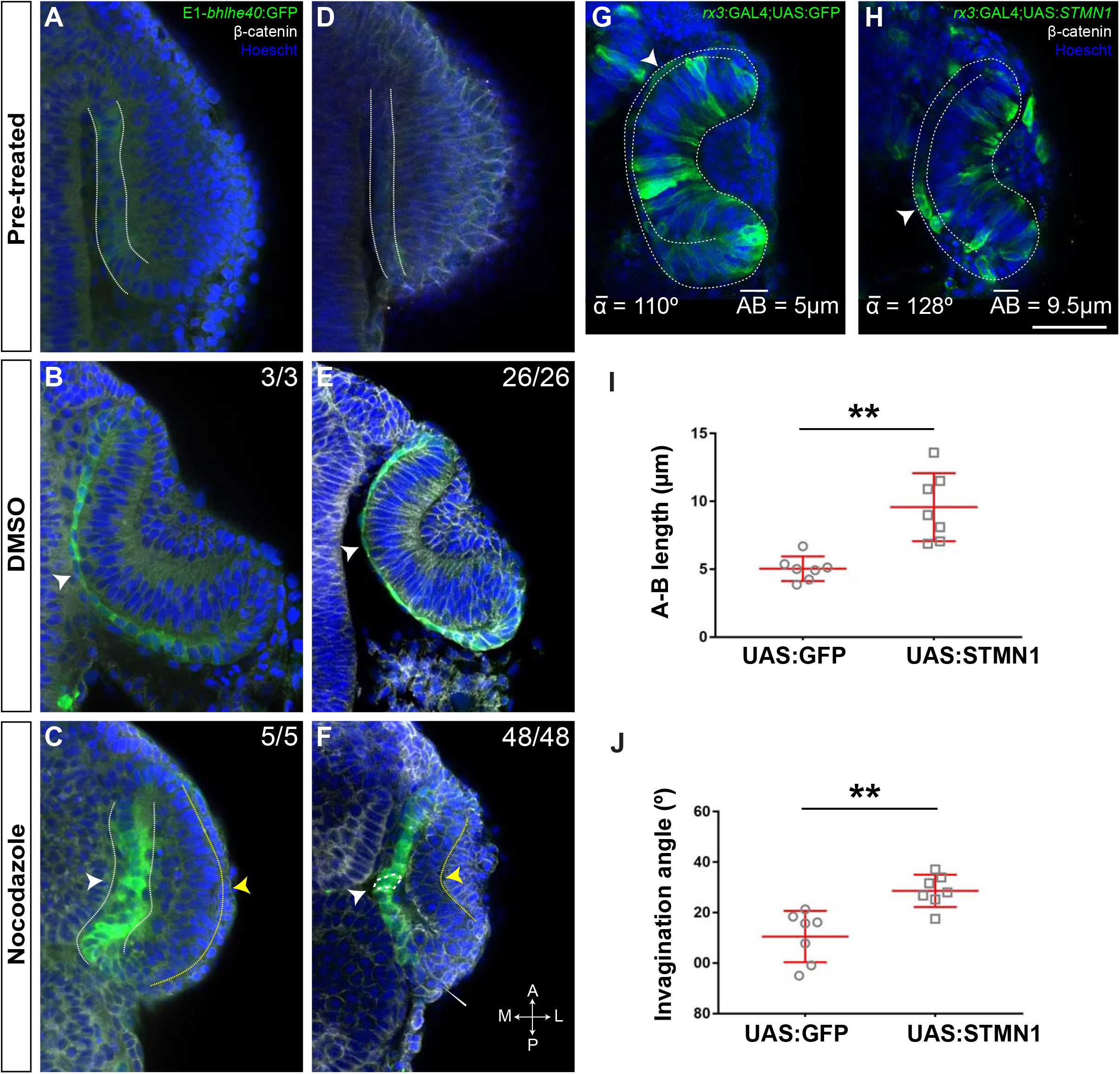
Microtubule dynamics is required for RPE cell flattening and OV folding. **A-F)** Confocal images of dorsally viewed Tg(E1-*bhlhe40*:GFP) embryos before (16 hpf, A; 17 hpf D) and 2.5 hr after incubation (18.5 hpf B, C; 19.5 hpf E,F) with either DMSO (B, E) or Nocodazole (C, F). Embryos were immunostained for GFP (green, A-F), β-catenin (white, D-F) and counterstained with Hoechst (blue). Note that the OC forms and the RPE flattens (white arrowhead in B, E) normally in DMSO treated embryos. RPE cells retain a columnar-like morphology in the presence of Nocodazole (white arrowhead in C, F). In embryos treated at earlier stage, the NR seems to bend outward (yellow arrowhead in C), whereas some folding occurs when the embryos are treated at later stages (yellow arrowhead in F), although cells seems to accumulate at the hinge (thin white arrowhead, F). The number of embryos analyzed and showing the illustrated phenotype is indicated on the top right corner of each panel. **G, H**) Confocal images of dorsally viewed *rx3*:GAL4;UAS;RFP embryos injected with the UAS:GFP (G, n=7) or UAS:*STMN1* (H, n=7) at one cell stage and fixed at 24 hpf. Embryos were labeled with anti-GFP (green) and counterstained with Hoechst (blue). Note that *STMN1* overexpression but not GFP prevents RPE cell flattening and cells retain a cuboidal like shape (white arrowheads in G-H). The average invagination angle and mean length of the A-B axis are indicated in the bottom left and right angles, respectively. **I, J**) The graphs show the length of the A-B axis (I) and the invagination angle (J) in embryos overexpressing GFP or *STMN1*. Mean ± SD. ** p<0.01). Scale bar: 50 μm.

The differential effect of nocodazole in the two layers of the OV further supports that RPE morphogenesis takes place rather independently from the adjacent NR layer, although whole embryo treatment did not unquestionably prove it. Unfortunately, we were unable to uncouple the effect of microtubule alterations in the two OV layers with a localized drug interference. We thus resorted to inject Tg(*rx3*:GAL4) embryos with either UAS:GFP or UAS:*STMN1*, encoding stathmin 1, a key regulator of microtubule depolymerization (44). We reasoned that, although *rx3* will drive transgene expression in both NR and RPE progenitors, the random and sparse expression occurring in F0, would be sufficient to separate the effect in the two tissues. RPE cells expressing *STMN1* – and notably also those nearby – retained a cuboidal-like shape with an abnormally increased apico-basal axis as compared to UAS:GFP expressing cells-, whereas cells in the inner OV layer appeared to undergo basal constriction (Fig. 5G-I). This phenotype correlated with poor OV invagination (Fig. 5J), supporting the notion that proper RPE flattening contributes to OV folding.

All in all, these data indicate that the RPE actively participates in OV folding by undergoing a tissue autonomous stretching driven by cell cytoskeletal rearrangements.

### Differential requirement of cell proliferation in zebrafish vs amniotes RPE development

Our finding that the zebrafish RPE largely grows through individual cell flattening agrees with the observation that zebrafish RPE cells barely proliferate during OV folding (22). This however differs from reports in mouse embryos, in which RPE proliferation seems a requirement for OV folding (27) and suggests the existence of species-specific modes of early RPE growth. We hypothesized that these modes may be related to the speed of embryonic development with final consequences on the epithelial characteristic of the RPE. To test this possibility, we compared proliferation rate and apico-basal length of the zebrafish RPE with those of the medaka, chick, mouse and human embryos at equivalent OV/OC stages. In these species, Otx2 and N-cadherin immunostaining was used to identify the RPE domain (23, 45) and the cell shape (Fig. S4), respectively.

BrdU incorporation in Tg(E1-*bhlhe40*:GFP) embryos from early OV (17 hpf) to late OC stages (48 hpf) showed a marked reduction of cell proliferation in the OV outer layer (Fig. 6A-B), very much in line with the report that only 2% of the outer layer cells undergo mitosis in this period (22). At the earlier stage (17 hpf), BrdU positive cells were scattered across the RPE with no easily identifiable geometry and accounted for 49% of the total RPE cells. This fraction dropped to about 20% at 19 hpf, when cells are flat, and then to 12% at 48 hpf (Fig. 6B) when the epithelium is maturing. Statistical analysis showed significant differences between 17 and 20 hpf (Mann-Whitney U test: z=-2.619, p<0.01, mean rank for 17 hpf is 8 and for 19 hpf is 3) and a clear correlation between proliferation rate and developmental stage (Kruskal-Wallis test: χ^2^(df=7, n=40)=31,864; p<0.001). During this period, the apico-basal axis of individual RPE cells flattened reaching a length of 3 μm at 22-23 hpf. Thus, acquisition of RPE identity, cell shape changes and OV folding are associated with a progressive reduction of cell proliferation in the OV outer layer. To determine if this reduction is a prerequisite for RPE flattening we injected a UAS*:ccnd1* construct into the *rx3*:GAL4 transgenic line (46), thus forcing the expression of a key regulator of the cell cycle (*ccnd1*) in a few OV progenitors (Fig 6C-H). Again, we reasoned that sparse expression occurring in F0 would be sufficient to separate the effect in the two tissues. Forced *ccnd1* expression maintained RPE progenitors in a proliferative state as detected by the presence of BrdU in *ccnd1* expressing cells (Fig. 6C,D). The small increase in BrdU/*ccnd1* positive RPE cells (52.5% *rx3*:GAL4:UAS-*ccnd1* vs 47.5%, *rx3*:GAL4:UAS-GFP; Fig. 6C-D) interfered with RPE specification and flattening (Fig. 6E-H). Indeed, *bhlhe40* mRNA distribution remained dorsally restricted in UAS:*ccnd1* injected embryos as compared to UAS:*GFP* injected controls (Fig. 6G, H), suggesting that in zebrafish acquisition of RPE properties requires fast cell cycle exiting.

**Figure 6.**
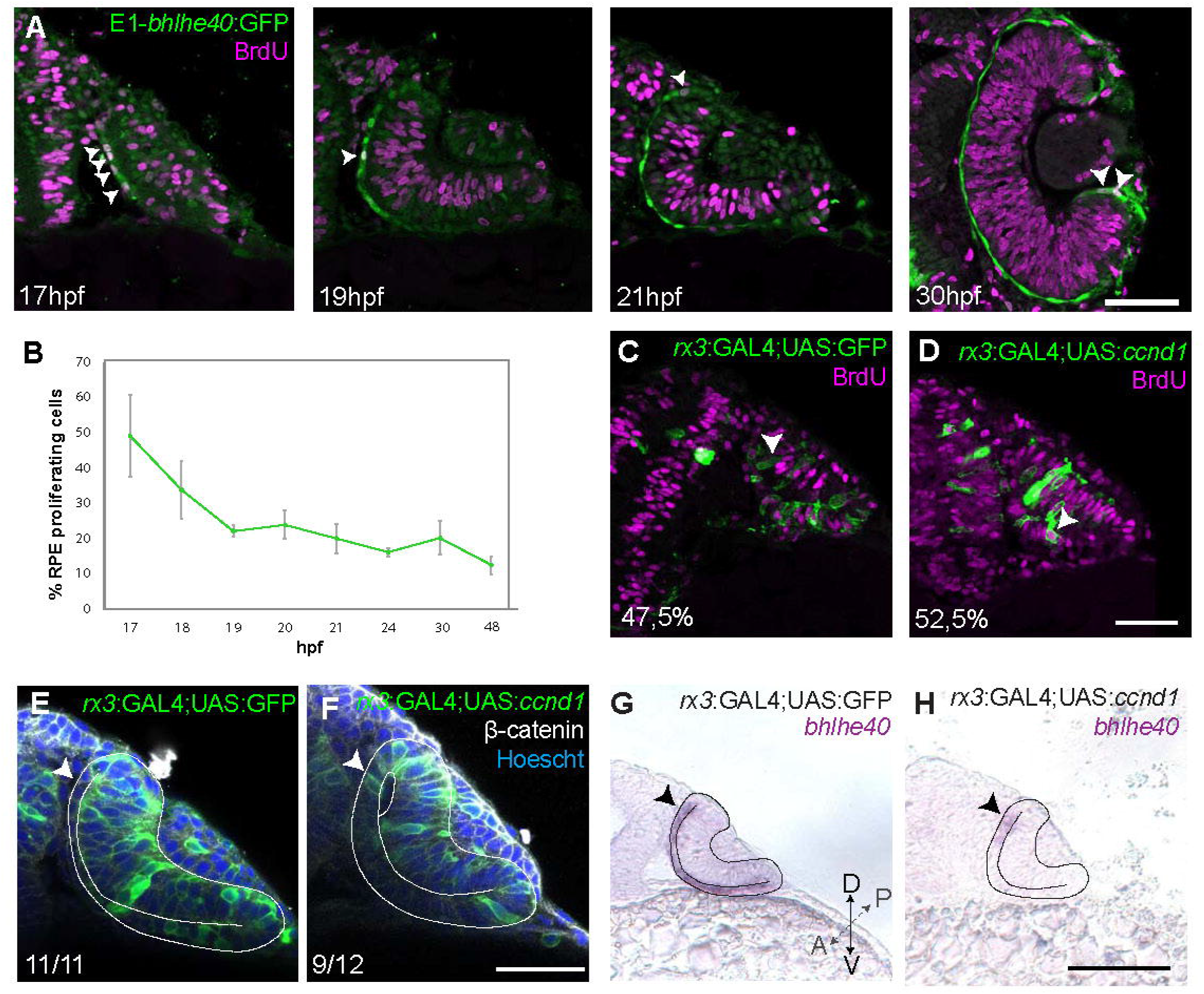
Zebrafish RPE flattening is associated with an abrupt decrease of cell proliferation. **(A)** Confocal images of dorsally viewed Tg(E1-*bhlhe40*:GFP) embryos exposed to BrdU at different developmental stages as indicated in the panel and immunostained for BrdU (magenta) and GFP (green). **B)** Percentage of RPE proliferating cells (BrdU+/total Hoechst +) in 17-48 hpf Tg(E1-*bhlhe40*:GFP) embryos. Mean ± SD; n=5 embryos per stage. **C, F)** Confocal images of dorsally viewed Tg(*rx3*:GAL4;UAS;RFP) embryos injected with either UAS:GFP (C,E) or UAS:*ccnd1* (D,F) and immunostained for BrdU (magenta) and GFP (green) (C-D) or GFP (green), β-catenin (white) and counterstained with Hoechst (blue) (E-F). Note that *ccnd1* over-expression in the RPE maintains cells with an elongated/cuboidal shape (white arrowhead in F) as compared to GFP expressing cells (white arrowhead in E). **G-H)** Dorsal views of Tg(*rx3*:GAL4;UAS;RFP) embryos injected with either UAS:GFP (G) or UAS:*ccnd1* (H) *and* hybridized with *bhlhe40* specific probe. Note that *ccnd1* overexpression prevents the expansion of *bhlhe40* expression which remains confined to the dorsal region of the outer layer (black arrowheads). Scale bars: 50 μm in C-F and 100 μm in G-H.

OC morphogenesis in the teleost medaka fish occurs with a choreography comparable to that of the zebrafish (17, 18) but the medaka fish RPE does not adopt an extreme squamous morphology (Fig 7A). Notably, medaka fish develop slower than zebrafish embryos, so that, from first appearance, their OVs take about eight hours more to reach a fully developed OC (26 vs 18 hr) (47), a time compatible with an additional round of cell division. Consistent with this idea, BrdU incorporation in st18 to 22-23 medaka embryos showed that about 70% of the cells in the OV outer layer were actively cycling and this proportion dropped to about 48% at OC stage (Fig. 7A,E) with a slightly less evident decrease of the average apico-basal axis (st18: 21.3μm vs st23 3.5μm; Fig 7A) as compared to the changes observed in zebrafish. An equivalent analysis in chick and mouse embryos showed similar results. In these species OV conversion into an OC takes about 27 and 48 hr, respectively. During this period a similar and almost constant proportion of RPE progenitors incorporated BrdU (Fig. 7B,C,E), including when cells acquired the expression of the RPE differentiation marker Otx2 (Fig. S4A). Furthermore, RPE cells only roughly halved their apico-basal axis (chick: HH12: 30.1μm vs HH18: 15.8μm; mouse: E9.5: 23.7μm vs E11.5 13μm Fig. 7B,C), suggesting that in slower developing species, proliferation but not stretching accounts for RPE surface increase. To corroborate this idea, we next analyzed human embryos.

**Figure 7.**
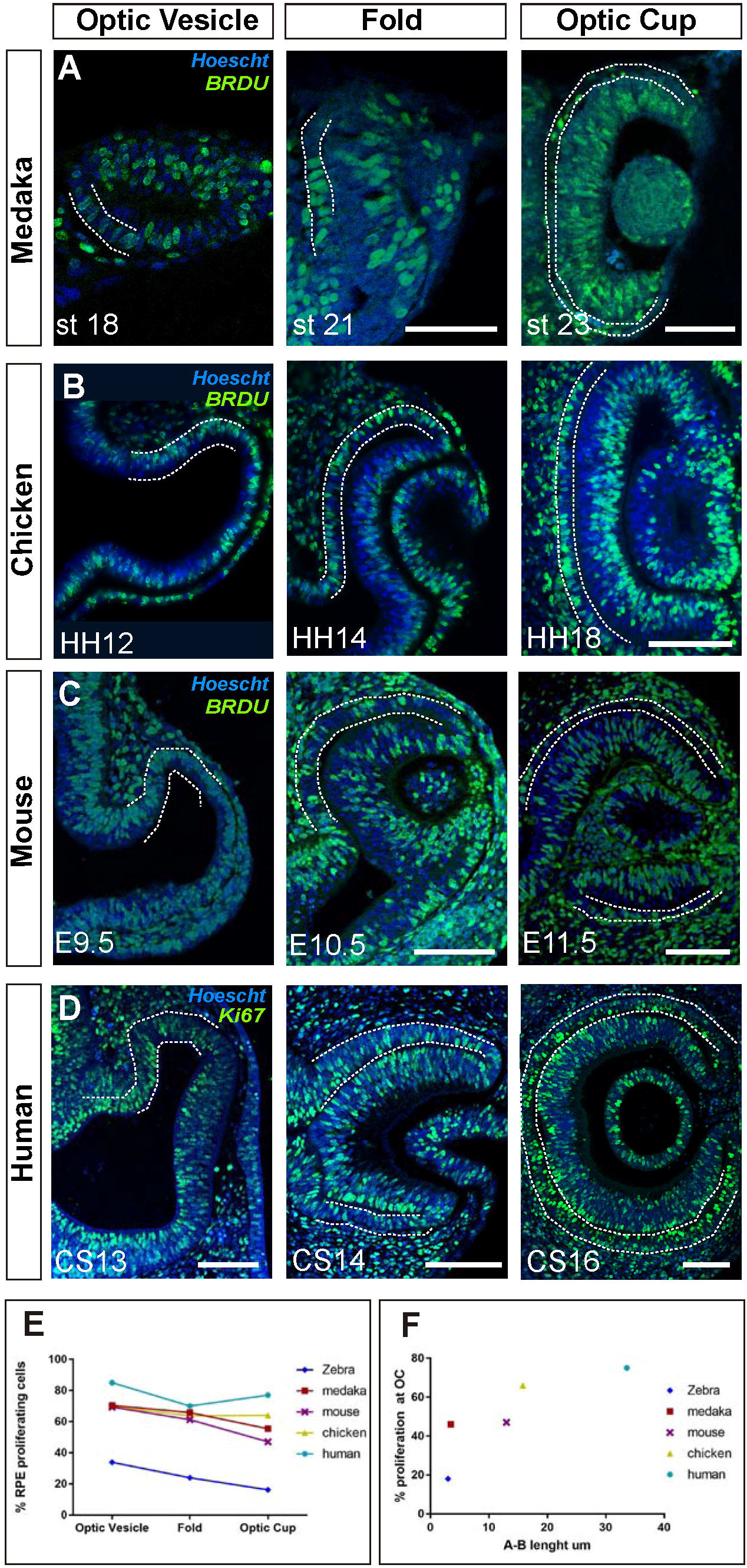
Proliferation accounts for RPE surface increase during amniotes OV folding. **A-C**) Confocal images of frontal sections from medaka, chick and mouse embryos exposed to BrdU at equivalent stages of OV folding into OC, as indicated in the panels. Sections were immunostained for BrdU (green) and counterstained with Hoeschst (blue). In all panels the prospective RPE has been highlighted by dotted white lines on the basis of the Otx2 immunostaining illustrated in Fig S4. D) Confocal images of horizontal sections from human embryos at equivalent stages of OV folding into OC. Sections were immunostained for Ki67 (green) and counterstained with Hoeschst (blue). **E)** Percentage of RPE proliferating cells (BrdU+/total Hoechst+) in the analyzed period and compared to those reported in Fig 6B for zebrafish. Data represent mean ± SD; the number of embryos analyzed for each stage varied between 3 to 10. **F)** Relationship between proliferation rate and apico-basal axis length at OC stage in the different species. Note that there is a positive correlation between the two parameters. Scale bars: 50 μm in A and 100 μm in B-D.

The human eye primordium is first visible at about four-five weeks of gestation corresponding to Carnegie stage (CS)13 (48). A fully formed OC is reached only roughly 10 days after, at CS16 (48). Immunostaining of paraffin sections from CS13 to CS16 embryos with antibodies against Ki67, a marker of the active phases of the cell cycle, demonstrated that the large majority of prospective RPE cells undergo a marked proliferation during the transition from OV to OC (Fig. 7D). Owing to the difficulties in obtaining early human embryonic samples, the percent of proliferating cells could only be estimated, showing that in the OTX2-positive domain (Fig. S4B), Ki67-positive RPE cells represented about 85% to 75% of the total between CS13 and 16. During this period, the prospective RPE layer always appeared as a rather thick pseudostratified epithelium with an organization resembling that of the NR composed of densely packed and elongated neuroepithelial cells (Fig. 7D; S4B). During the formation of the OC, the RPE neuroepithelium only slightly flattened (apico-basal thickness: CS13: 45μm vs CS16: 33.6μm), far from reaching the cuboidal appearance seen at postnatal ages (Fig S4B).

Collectively these data indicate that, in the absence of sufficient time for cell proliferation, flattening is an efficient solution adopted by zebrafish RPE cells to enlarge the whole tissue to the extent needed for OV folding. In other vertebrates, in which slower development allows for more rounds of cell division, the RPE grows in a conventional proliferation-based mode that correlates with a less evident flattening of RPE cells (Fig. 7F).

## Discussion

The cup-shape of the vertebrate eye is thought to optimize vision (49). This shape is acquired very early in development as the result of specification and morphogenetic events, during which the NR and the RPE arise. Studies in teleosts (zebrafish and medaka) together with mammalian organoid cultures have recently demonstrated a fundamental contribution of NR progenitors in driving the acquisition of this cup shape (2, 9). The role of the RPE progenitors in this process has instead not been properly clarified. In this study, we have filled this gap and analyzed the folding of the zebrafish OV from the RPE perspective. This analysis has been possible thanks to the generation of new RPE reporter line (Tg(E1-*bhlhe40*:GFP), in which GFP expression appears in the domain fated to originate the RPE. Following the cells arising from this domain, we show that RPE surface expansion is an active and tissue autonomous process that is needed for the proper folding of the OV. This expansion largely occurs by extreme cell flattening with little contribution of cell proliferation, a mechanism that sets zebrafish RPE morphogenesis apart from that of other analyzed vertebrate species, in which proliferation accounts for RPE growth.

Our analysis together with a previous report (22) shows that the onset *bhlhe40* expression coincides spatially and temporally with the onset of zebrafish RPE specification. Thus, the Tg(E1-*bhlhe40*:GFP) line serves as an early tissue specific marker that even precedes the appearance of previously accepted *Otx* or *Mitf* tissue specifiers, as confirmed in a parallel transcriptomic analysis (8). *Bhlhe40* expression in the RPE is conserved at least in mouse and humans (8,50,51), suggesting a possible relevant function in this tissue. However, its CRISP/Cas9 inactivation, alone or in conjunction with that of the related *bhlhe41, mitfa* and *mitfb*, had no evident consequences on zebrafish RPE development, at least in our hands (data not shown). One possible reason for the absence of an evident RPE phenotype is functional redundancy with other untested members of the large family of the BHLH transcription factors or that the genes might have only later functions as reported (52). However, we favor the alternative possibility that zebrafish RPE specification may occur “en bloc”, making the inactivation of one or two genes insufficient to perturb fate acquisition. Such a mechanism is expected to provide robustness to a process that takes place in just few hours and finds support in present and past findings (8, 22).

Indeed, we and others (22) have shown that, by the time the OV start to bend, the large majority of RPE cells have already left the cell cycle and have acquired a differentiated squamous morphology by undergoing a marked surface enlargement in the medio-lateral direction, while reducing the apico-basal axis, with the net result of an overall modest volume increase. Transcriptomic analysis shows that during this same lag of time, RPE cells repress genes characteristic of 16 hpf OV progenitors, such as *vsx1*, and acquire the expression of RPE specific genes. These includes blocks of transcription factors, such as known RPE specifiers (i.e. *otx, mitf*) and regulators of epidermal specification (i.e. *tfap* family members, known regulator of keratin gene expression (53)) as well as several cytoskeletal components, most prominently a large number of keratins and other desmosomal components found in squamous epithelia (8). Thus, in just few hours (from 16 to 18 hpf) RPE cells acquire the molecular machinery required for their conversion from a neuroepithelial to a squamous epithelium. Our study shows that this conversion relays on a cell/tissue autonomous cytoskeletal reorganization without the influence of the morphogenetic events occurring in the nearby NR. Indeed, local interference with actomyosin or microtubule dynamics is sufficient to retain RPE cells into a cuboidal or neuroepithelial configuration, respectively, without affecting their specification. In contrast, localized interference with NR bending has no effect on RPE flattening.

The extreme flattening of the zebrafish RPE cells makes the resolution of their cytoskeletal components difficult with *in vivo* confocal microscopy, hampering the complete understanding of how the actomyosin cytoskeleton promotes the acquisition of a squamous configuration. In other contexts, a flat morphology is associated with the presence of actomyosin stress fibers that compress the nucleus (36, 37). Myosin II is essential for this compressive role and its inhibition with blebbistatin causes the loss of the flat morphology (36, 37), as we have observed in blebbistatin and azoblebbistatin treated embryos. It is thus possible that a similar nuclear compression may occur in the RPE cells as they flatten, although we were unable to detect stress fibers around the nucleus, likely due to plasma membrane proximity. Remodeling of the microtubular cytoskeleton seems to further aid RPE cell flattening. Microtubules change their orientation during RPE morphogenesis, from being aligned along the apico-basal axis of the cells at the onset of RPE morphogenesis, to becoming aligned with the planar axis in squamous RPE cells. A similar process has been described during the morphogenesis of the *Drosophila* amnioserosa (42), in which cells also change from a columnar to a squamous morphology. In these cells, actin accumulation at the apical edge offers resistance to the elongation of microtubules, which thus bend, leading to a 90° rotation of all subcellular components. This rotation is accompanied by a myosin-dependent remodeling of the adherens junctions (42), a process that may also take place during RPE flattening.

Although additional studies are needed to clarify the precise dynamics of the cytoskeletal reorganization underlying RPE differentiation, our study demonstrates that cytoskeletal dynamics occurs in a tissue autonomous manner. In contrast to other studies (11, 13), we have used a photoactivable version of blebbistatin that has allowed us to determine the individual contribution of the NR and RPE to OV folding. As a drawback, this approach allows to activate the drug only in relatively small patches of tissue. It was thus rather remarkable to observe that failure of RPE flattening in small regions was sufficient to decrease OV folding. This suggests that RPE stretching represents an additional and relevant mechanical force that, together with retinal basal constriction and rim involution, contributes to zebrafish eye morphogenesis (Fig 8A). This flattening and stretching together with a substantial expression of keratins (8) may confer a particular mechanical strength to the zebrafish RPE, which may constrain the NR and, at the same time, may favor rim involution (17). The latter possibility is supported by the observation that inner layer cells seems to accumulate at the hinge in the absence of RPE flattening. These marked morphogenetic rearrangements can thus be seen as an efficient solution adopted in fast developing species to make eye morphogenesis feasible in a period does not to allow for proliferation-based tissue growth.

**Figure 8.**
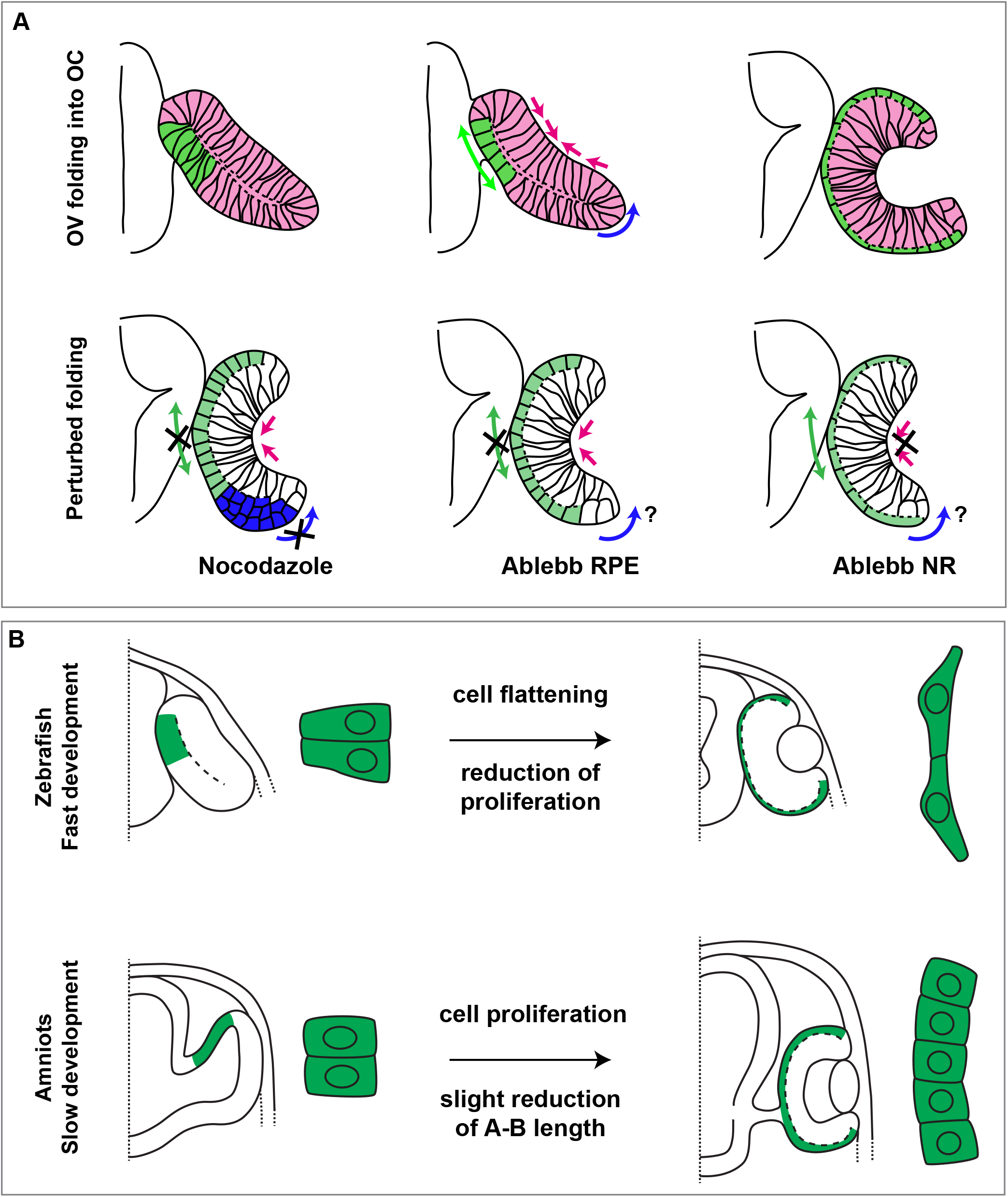
Summary of species-specific modes of RPE differentiation and its contribution to OV folding. **A)** The drawing on the top represent the dynamic of OV folding into an OC. Green double arrow indicated RPE flattening, blue arrow rim involution whereas pink arrows indicate retinal basal constriction. Bottom row summarize the alterations in OV folding observed after localized interference with RPE and NR cytoskeleton. **B)** Schematic representation of the differential mechanisms by which the RPE in zebrafish (upper row) and in amniotes (lower row) expands its surface during OV folding morphogenesis. In zebrafish, the RPE enlarges its surface by cell stretching; in amniotes, including in humans, the RPE instead expands by cell proliferation with a less pronounced need of cell flattening.

The perhaps obvious question is whether similar morphogenetic rearrangements are needed in other vertebrates to form the remarkably conserved cup shape of the eye. So far, rim involution has been reported only in teleost species where it may represent a fast mode of increasing the surface of the inner layer of the OV, thus favoring its bending (13,17,18); well in agreement with previous data showing that between 16 to 27hpf the outer layer of the OV decrease its cell number from about 587 to 432, whereas the inner layer increases its numbers in a way that cannot be explained solely by proliferation (16). In other species, this cell displacement may not be needed as the layer can grow by cell division. In a similar way, we have shown here that in slower developing species, RPE cells maintain a higher proliferation rate that contributes substantially to the increase of RPE surface while undergoing less marked changes in cell shape (Fig. 8B). This correlation is visible in medaka, despite its relative evolutionary proximity to zebrafish (47), and it is maximal in human embryos. Indeed, in humans, the RPE layer is composed of cells with a neuroepithelial appearance and a high proliferation rate, despite the expression of OTX2, considered a tissue specifier. Thus in mammals, full commitment of the OV outer layer to an RPE identity may occur over a prolonged period of time and not “en bloc” as in zebrafish, as suggested by comparing RNA-seq data from RPE cells from human CS13-16 embryos (51) with those from equivalent stages in zebrafish (8). Human RPE cells from CS13-16 embryos are still enriched in the expression of proliferation associated genes (51) but not of those typical of squamous epithelia as in zebrafish (8). A slow acquisition of RPE identity may also explain why inactivation of genes such as *Otx2, Mitf* or *Yap* causes the RPE layer to adopt NR characteristic (23,25,54), whereas this feature that has never been reported after equivalent manipulations zebrafish (55, 56), or why FGF8 can push the amniote but not the zebrafish RPE layer to acquire a NR identity (57). As a reflection of this slower differentiation in amniotes, RPE cells can largely retain their neuroepithelial morphology and adopt a final cuboidal-but not squamous-appearance at a slower and species-specific pace.

We thus propose that RPE cell stretching versus cell addition are different solutions adopted by species with different rates of development to reach a common goal: reaching the appropriate equilibrium between the surface of the RPE and of the NR. Indeed, the present study together with previous observations (27) and *in silico* models (15, 58) support that this equilibrium is a prerequisite for proper OV folding.

## Material and Methods

### Animals

Adult zebrafish (*Danio rerio*) were maintained under standard conditions at 28°C on 14 h-light/10 h-dark cycles. AB/Tübingen strain was used to generate the transgenic lines and as control wild type. Embryos and larvae were kept in E3 medium (5 mM NaCl, 0.17 mM KCl, 0.33 mM CaCl2, 0.33 mM MgSO4) supplemented with Methylene Blue (Sigma) at 28°C and staged according to somite number and morphology (59). The lines Tg(E1-*bhlhe40*:GFP) and Tg(*rx3*:Gal4;UAS:RFP) (46) lines were maintained in the same conditions and crossed to generate the Tg(E1-*bhlhe40*:GFP;*rx3*:GAL4;UAS;RFP) line. Wild-type medaka fish (*Oryzias latipes*) of the cab strain were maintained at 28°C on a 14/10-hour light/dark cycle. Embryos were staged as described (60). Fertilized chick embryos (Santa Isabel Farm, Cordoba-Spain) were incubated at 38°C in a humidified rotating incubator until the desired stage. Embryos were inspected for normal development and staged according to (61). Wild-type BALB/c mice were in pathogen–free conditions at the CBMSO animal facilities, following current national and European guidelines (Directive 2010/63/EU). The day of the appearance of the vaginal plug was considered as embryonic day (E)0.5. All experimental procedures were approved by the CBMSO and Comunidad Autónoma de Madrid ethical committees.

### Human tissue

Paraffin sections of human embryonic eye primordia were obtained from Human Developmental Biology Resource (http://www.hdbr.org/) through the University of Newcastle upon Tyne and the University College London under the grant # 200498) and according to ethics approval from all the involved institutions (UCL; 18/LO/0822 - IRAS project ID: 244325; UNuT, 18/NE/0290 - IRAS project ID: 250012; CSIC-CBMSO, 003/2020). Sections corresponded to samples Carnegie Stage (CS) 13, 14, 15 and 16. CS staging allowed to determine the age of embryo as days post ovulation based on morphological landmarks (62).

### Generation of the Tg(E1-bhlhe40:GFP) line

Predictive enhancer and promoter epigenetic marks (30) were used to identify different potential regulatory elements of the *bhlhe40* gene (Fig. 1B). Each region was amplified by PCR with specific primers (Table S1) and cloned using the pCR™8/GW/TOPO^®^ TA Cloning^®^ Kit (Invitrogen). Plasmids were checked for enhancer insertion and the Gateway™ LR Clonase™ Enzyme Mix (Invitrogen) was used for recombination with the ZED vector (31). The resulting constructs were injected together with Tol2 mRNA to generate the corresponding transgenic embryos, which were screened using a transgenesis efficiency marker present in the ZED vector (cardiac actin promoter:RFP). Positive larvae were grown to adulthood (F0) and then individually outcrossed with wild type partners to identify founders. Founders were analysed using confocal microscopy. One of the lines corresponding to the enhancer E1 was finally selected and used for subsequent studies.

### Gal4-UAS-mediated expression

The *UAS:STMN1* and *UAS:ccnd1* constructs were generated from the bi-directional UAS:GFP vector (63). Both genes were amplified by PCR using specific primers (Table S1) flanked by StuI restriction sites and the Expand™ High Fidelity PCR System. STMN1 human gene was amplified from the pQTEV-STMN1 (Addgene #31326) and *ccnd1* zebrafish gene isolated from 24 hpf cDNA. Both PCR products were digested with StuI (Takara) and cloned into the pCS2 vector and thereafter isolated together with the polyA sequence of the vector by digestion with HindIII and SacII (Takara) and subcloned into the UAS:GFP plasmid. The generated plasmids (30 pg) were injected into the Tg(rx3:Gal4;UAS:RFP) (46) line, together with Tol2 mRNA (50 pg) to increase efficiency.

### Embryos microinjection and drug treatments

Embryos at one cell stage were injected using a Narishige microinjector and glass needles prepared by horizontally pulling standard capillaries (filament, 1.0mm, World Precision Instruments) with aP-97 Flaming/Brown Micropipette Puller (Sutter Instrument Company). A total of 30 pg for DNA and between 50 and 100 pg for mRNA in 1 nl volume were injected in the embryos in the cell or the yolk, respectively. Drug treatments were performed on manually dechorionated embryos at the desired developmental stage in E3 medium. The following compounds were used: Blebbistatin (100 μM for 2.5 hr; Calbiochem); Paranitroblebbistatin (20 μM; Optopharma) Azidobblebistatin (5 μM for 15min before photoactivation; Optopharma) and Nocodazole (10 ng/μl for 2.5 hr; Sigma)

### In vitro transcription

The pCS2:Kaede, pCS2:EB3.GFP and pCS2:H2B-RFP constructs were linearized and transcribed using the mMessage mMachine™ SP6 transcription kit (Invitrogen), following manufacturer’s instructions. After transcription mRNAs were purified using the NucleoSpin^®^ RNA Clean-up kit (Machery Nagel).

### In situ hybridization (ISH)

*otx1* (previously known as *otx1b*) and *mitfa* probes were gifts from Prof. Steve Wilson (UCL, London UK). The *bhlhe40* probe was generated by PCR from 24 hpf cDNA with specific primers (Table S1) using the Expand™ High Fidelity PCR System. Reverse primers included the T3 promoter sequence to *in vitro* transcribe the PCR product. *In vitro* transcription was performed using T3 RNA polymerase and DIG RNA labelling Mix (Roche) following manufacturer’s instructions. Transcription products were precipitated with LiCl 0.4M and 3 volumes of ethanol 100% overnight at −20°C. Samples were centrifuged at 4°C and 12000 g for 30 min, washed with ethanol 70% and resuspended in 15 μl of RNAse free water and 15 μl Ultra-Pure Formamide (Panreac). ISH were performed as described (64).

### BrdU incorporation assays

BrdU (5-Bromo-2’-deoxyuridine, Roche) was resuspended in DMSO (Sigma) to generate stocks of 50 mg/ml that were kept at −20°C. For Tg(E1-*bhlhe40*:GFP) zebra- and wild type medaka fish groups of 15 embryos of stages comprised between 16 ss and 48 hpf were dechorionated and placed in BrdU solution (5 mg/ml in E3 medium) for 30 min on ice and then washed with fresh E3 medium. Embryos were let recover at 28°C for 10 min before fixation in PFA 4% overnight at 4°C. For analysis in chick, BrdU (50 mg/egg) was added to each embryo 30 min before fixation. For analysis in mouse, pregnant dams were injected intraperitoneally with BrdU (50 μg/g), sacrificed 1hr later and fixed. Chick and mouse embryos were immersion fixed in 4% paraformaldehyde in 0.1 M phosphate buffer, pH 7 at 4°C for 4hr and then washed in PBS and cryoprotected in 15% and 30% saccharose in 0.1 M phosphate buffer. All embryos were cryo-sectioned and the sections hydrated with PBS 1X during 5 min and incubated in HCl during 40 minutes at 37°C. After HCl treatment, sections were rinsed with PBS 1X ten times, and then processed for immunofluorescence as described below. The percentage of RPE proliferating progenitors was determined as the proportion of BrdU positive cells over the total number of GFP (for E1-*bhlhe40*:GFP) or Otx2/Hoechst (medakafish, chick, mouse embryos) positive cells in the RPE layer in each section. A minimum of three embryos and sections per embryo were counted (both eyes).

### Immunofluorescence

Zebrafish embryos at the corresponding stage for each experiment were fixed with 4% (w/vol) paraformaldehyde (PFA, Merck) in 0.1 M phosphate buffer overnight at 4°C. Whole-mount immunofluorescence was performed as described (64). Alternatively, embryos were incubated in 15% sucrose - PBS overnight at 4°C, embedded in 7.5% gelatine (Sigma) 15% sucrose (Merck), frozen in isopentane (PanReac) between −30 and −40°C and kept at −80°C. Cryo-sectioning was performed with a cryostat (Leica CM 1950) at 20 μm thickness and dried overnight at RT. Chick and mouse embryos were collected, fixed 4% paraformaldehyde, equilibrated in sucrose and cryo-sectioned as above. Paraffin sections of human embryonic tissue were de-paraffinized, washed in PBS, processed for antigen retrieval (10mM citrate buffer, pH6, for 5min at 110°C in a boiling chamber, Biocaremedical) and subsequently processed together with all other samples for immunofluorescence.

Immunostaining was performed as described (64) using the following primary antibodies: mouse anti-BrdU (1:200; Becton-Dickinson); chick anti-GFP (1:2000; Abcam); mouse anti-βcatenin (1:400, BD transduction Laboratories); mouse anti-ZO-1 (1:400, Invitrogen); rabbit anti-laminin (1:200, Sigma); rabbit anti-Otx2 antibodies (1:1000; Abcam) rabbit anti-Ki67 (1:500, Abcam). The used secondary antibodies were conjugated with Alexa 488, Alexa-594 or Alexa-647 (1:500; Thermo Fisher). Sections were counterstained with Hoechst (Invitrogen), mounted in Mowiol and analysed by conventional and confocal microscopy.

### Kaede photoconversion

Wild type embryos were injected with Kaede mRNA. Embryos at 15 hpf with homogeneous green fluorescence were selected, mounted and visualized under the Nikon AR1+ Confocal Microscope using a 20X/0.75 Plan-Apochromat objective. A region of interest (ROI) was drawn in the outer layer, corresponding to the putative position of the RPE progenitors, at a specific z position and irradiated with the 405 nm laser at 21% of power for 10 loops to switch Kaede emission from green to red fluorescence. Due to confocality, photoconversion occasionally extended further than the selected plane, so that the tissues present above or below (i.e. ectoderm) also underwent photo-conversion. After photo-conversion embryos were let develop up to approx. 30 hpf stage, fixed and analysed by confocal microscopy for red fluorescence distribution.

### Azidoblebbistatin photoactivation

Azido-blebbistatin (Ableb) (39) was photoactivated with a Zeiss LSM 780 Upright multiphoton FLIM system with a W Plan-Apochromat 20x/1,0 DIC M27 75 mm WD 1.8 mm dipping objective. For each eye a specific ROI was drawn including RPE cells identified by GFP fluorescence. ABleb was activated in the ROIs using 860 nm wavelength and 20 mW laser power (this corresponds to 9-14 μW/μm2 inside the ROI).

### Confocal imaging

Embryos were mounted with the appropriate orientation in 1.5% low melting point agarose (Conda) diluted in E3 medium (for in vivo recording) or PBS (for fixed samples). Images were acquired either with a Nikon A1R+ High Definition Resonant Scanning Confocal Microscope connected to an Inverted Eclipse Ti-E Microscope (20X/0.75 Plan-Apochromat, 40X/1.3 oil Plan-Fluor and 60X/1.4 oil Plan-Apocromat objectives) or with a Zeiss LSM710 Confocal Laser Scanning Microscope connected to a Vertical AxioImager M2 Microscope (40X/1.3 oil Plan-Apochromat, W N-Achroplan 20x/0.5, W Plan-Apochromat 40x/1.0 DIC VIS-IR).

### 3D reconstructions

3D movies (i.e. Movie S1-3) were generated from full stacks using the 3D project option in Fiji (65). RPE surface renderings were generated using Imaris (Bitplane), with a value of 6 in Surface Area Detail and 7 in Background Subtraction.

### Morphometric analysis

Unless otherwise specified, morphometric analysis of cells and tissues was performed using Matlab© (The Mathworks©, Natick, MA) using the XYZ coordinates of the processed images or Fiji (65). This analysis was performed using previously processed fluorescent images from movies of Tg(E1-*bhlhe40*:GFP; *rx3*:GAL4;UAS:RFP) or Tg(E1-*bhlhe40*:GFP) and H2B-RFP-injected embryos (Movie S2 and S3), from which the signal corresponding to the RPE or the whole OV/OC were isolated semi-manually with the help of Fiji macros and tools designed to select 3D structures. The RPE specific GFP signal was processed with a median filter. In the case of Movie S3, the background ramp for the GFP signal was neutralized in each frame via subtraction of a copy of itself after a gray-scale morphological operation (66, 67). For all Movies, the median intensity was thereafter established as the cutoff value for differentiating background and signal (i.e pixel with an intensity lower than the cutoff were set to zero) for all images that were in both movies. The signal derived from H2B was localized in cell nuclei, and therefore it was post-processed with a grayscale closing operation to fill empty spaces between nuclei. Morphometric analysis was performed in the resulting processed images. All values were calculated in microns by scaling the x,y,z coordinates according to the follow: (0.62μm x 0.62μm x 1.37μm) for movie S2 and (0.62μm x 0.62μm x 1.07μm) for movie S3. Volumes (μm^3^) were calculated as the number of voxels with a value higher than zero. RPE surface (μm^2^) was calculated applying a second order linear adjustment on the plane YZ corresponding to the plane of the OV/OC hinges with the fit function available in Matlab© (The Mathworks©, Natick, MA). RPE thickness (μm) was determined as the result of volume (μm^3^)/surface (μm2). Unfortunately, semi-manual RPE image extraction was not perfect, when GFP-signal associated to CMZ development arise. To account for this problem, the GFP signal for each frame was divided into seven equivalent blocks using the x,y coordinates from the z-projection of each frame. In this case, RPE volume and surface were calculated independently in each one of the regions up to 20 hpf, when the most anterior block (now corresponding to the arising CMZ) was discarded from the analysis. For the subsequent frames the two anterior most blocks were discarded (Fig. S1). The total OV/OC volume (μm^3^) was determined using the red fluorescence from the Tg(*rx3*:GAL4;UAS:RFP) embryos at 17-22hpf or the signal from H2B-RFP-injected embryos for stages 24 to 37 hpf calculating the convex hull structure of the retina (68) as the scaled number of voxels with a value higher than zero at each time point; the lens was considered as an empty space. Individual cell area was determined in cells located at a medial position of the OV for each cell type (progenitor, RPE and NR); cell contour was drawn using the segmented line tool in Fiji (65). Apico-basal (A-B) length (μm) of individual cells was estimated by manually tracing a line from the basal to the apical membrane in the z position in which the nucleus had its larger surface using the straight-line tool in Fiji (65). To account for possible developmental asynchrony when eyes from the same embryo were differentially treated (irradiated vs non irradiated), the A-B length of the irradiated eye was normalized with that of the non-irradiated eye. Values above 1 indicated less RPE cell flattening in experimental eyes. The invagination angle was determined as previously described (13) using manual drawing with the Fiji angle tool (65). The vertex of the angle was placed approximately in the center of the basal surface of the NR and the vectors were drawn up to the edges of the CMZ. Angles were measured in the z positions in which the irradiated RPE was maximally affected and compared to equivalent positions of control non irradiated eyes. Values were normalized with those of the contralateral non-treated eye, to account for possible asynchronies.

### Statistical analysis

All statistical analysis was performed with IBM SPSS Statistics Version 20.0. The method used is indicated in each case together with the sample size.

### Data Availability

Primary data and material generated in this study will be available upon request

## Acknowledgements

We wish to thanks Drs JR Martinez-Morales and E. Marti for critical reading of the manuscript. This work was supported by grants from the Spanish AEI (BFU2014-55918-P to FC and BFU2016-75412-R with FEDER support; RED2018-102553-T and PID2019-104186RB-I00 to PB), BBVA Foundation (N[16]_BBM_BAS_0078 to FC) and Fundación Ramon Areces-2016 (to PB). TMM and ML were supported by FPU (FPU14/02867) and FPI (BES-2015-073253) predoctoral contracts from the Spanish AEI, respectively. We also acknowledge a CBM Institutional grant from the Fundación Ramon Areces.

## Author contributions

TMM, FC and PB conceptualized and designed the research study. TMM performed most of the experiments, acquired and analysed the data. MLT performed bioinformatics quantifications reported in Fig. 3 and S1. NT, MJMB, MJC and PB generated data reported in Fig. 6C,D; Fig. 7 and Fig S4. TMM, FC and PB wrote the paper. All authors read and approved the manuscript.

## Supplementary Figure Legends

**Figure S1. RPE region selection from the GFP positive domain**. **A**) 3D reconstruction of the frames at 18, 19.7 and 21.5 hpf from the Movie S1. The analyzed GFP positive domain are depicted in green whereas those in turquoise have been excluded when calculating the RPE volume plotted in Fig. 3A. The *rx3* positive domain is depicted in pink. **B)** RPE volume bar plot. Light and dark green colors illustrate the total GFP positive volume of the central RPE and CMZ region respectively. The dark green region has been excluded to calculate the total volume.

**Figure S2. EB3-GFP dynamics during RPE cell remodeling. A-C)** Frames from representative movies S5-7 showing the orientation of microtubule dynamics in RPE cells with a neuroepithelial (A, continuous acquisition, n = 9), cuboidal (B, continuous acquisition, n = 10) and squamous conformation (C, continuous acquisition, n = 15). Scale bar 25 μm

**Figure S3. Nocodazole treatment does not alter RPE specification and polarity.** (A) DMSO and (B) Nocodazole-treated embryos (2.5 hr incubation from 17-19.5 hpf, as reported in Fig. 5) hybridized with *otx1* specific probe. (C) DMSO or (D) Nocodazole-treated embryos with *mitfa* specific probe. After Nocodazole treatment both mRNA are specifically detected in the RPE (black arrowheads) despite its morphogenesis is impaired. (E, E’) Wild type embryos labeled with laminin (green/basal), zo-1 (white/apical) and Hoechst (blue) after DMSO or (F, F’) Nocodazole treatment. In both cases apico-basal polarity is maintained (white and green arrowheads), even after depolymerizing microtubules when RPE cells do not flatten correctly. A-D images are dorsal views of flat mounted embryos. E-F’ images are frontal sections of one eye. Scale bars: 100 μm in A-D and 50 μm in E-F’

**Figure S4. Proliferation accounts for RPE surface increase during amniotes OV folding. A)** Confocal images of frontal sections from E9.5 and E10.5 mouse embryos coimmunostained for the RPE specific marker OTX2 (green) and actin (red). Sections were counterstained with Hoeschst (blue). **B)** Confocal images of horizontal sections from human embryos at CS15 and P5. The embryonic sections were immunostained for OTX2 (green) and N-cadherin (red) counterstained with Hoeschst (blue). Postnatal sections were immunostained for OTX2 (red). The distribution of OTX2 was used to define the prospective RPE region whereas actin/N-cadherin distribution to identify the apico-basal axis of the cells. Scale bars: 100 μm in A and left B panel; 30 μm in B right panel.

**Movie S1:** Dorsal view of the OV to OC transition visualized in a double Tg(E1-*bhlhe40*:GFP; *rx3*:GAL4;UAS:RFP embryo (related to Fig. 1, frame rate 1/5min).

**Movie S2:** Dorsal view of the OC folding visualized in a double Tg(E1-*bhlhe40*:GFP; *rx3*:GAL4;UAS:RFP embryo (related to Fig. 1, frame rate 1/5min).

**Movie S3:** Lateral view of the OC folding visualized in a double Tg(E1-bhlhe40:GFP; rx3:GAL4;UAS:RFP embryo (related to Fig. 1, frame rate 1/5min.

**Movie S4:** Lateral view of the OV to OC transition visualized in of an Tg(E1-*bhlhe40*:GFP) embryo injected with H2B-RFP mRNA (magenta) (related to Figure 3, frame rate 1/5min).

**Movie S5:** *eb3*GFP dynamics at 14 hpf. RPE cells still have a neuroepithelial organization (continuous acquisition, n = 9).

**Movie S6:** *eb3*GFP dynamics at 17 hpf. RPE cells have a cuboidal appearance (continuous acquisition, n = 10).

**Movie S7:** *eb3*GFP dynamics at 23 hpf. RPE cells have adopted a squamous conformation

