## Supplementary figures and images for "Stretching of the retinal pigment epithelium contributes to zebrafish optic cup morphogenesis"

### Fig S1

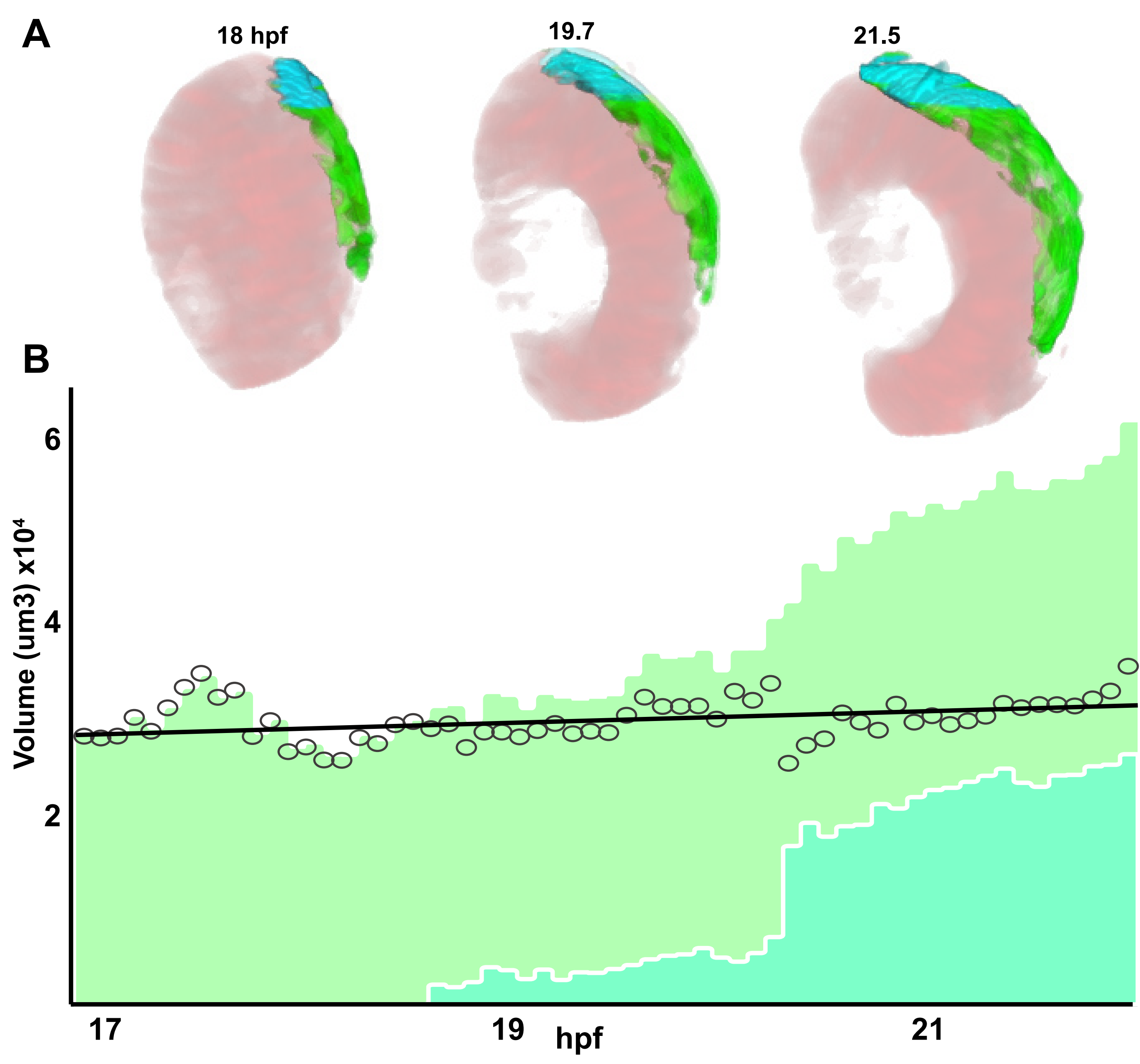

### Fig S2

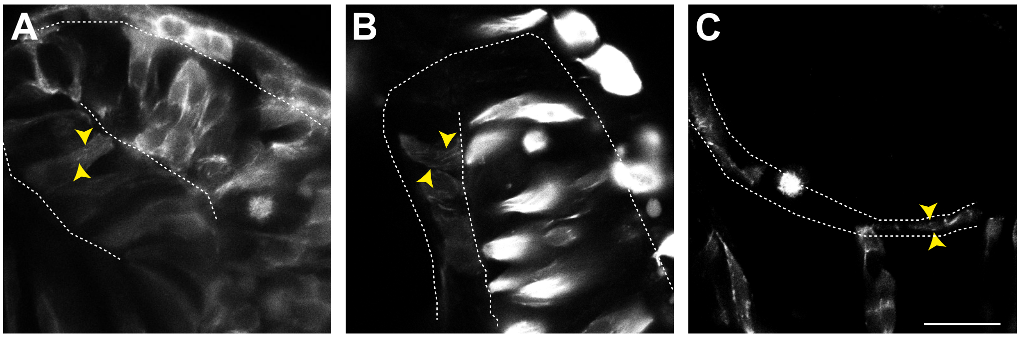

### Fig S3

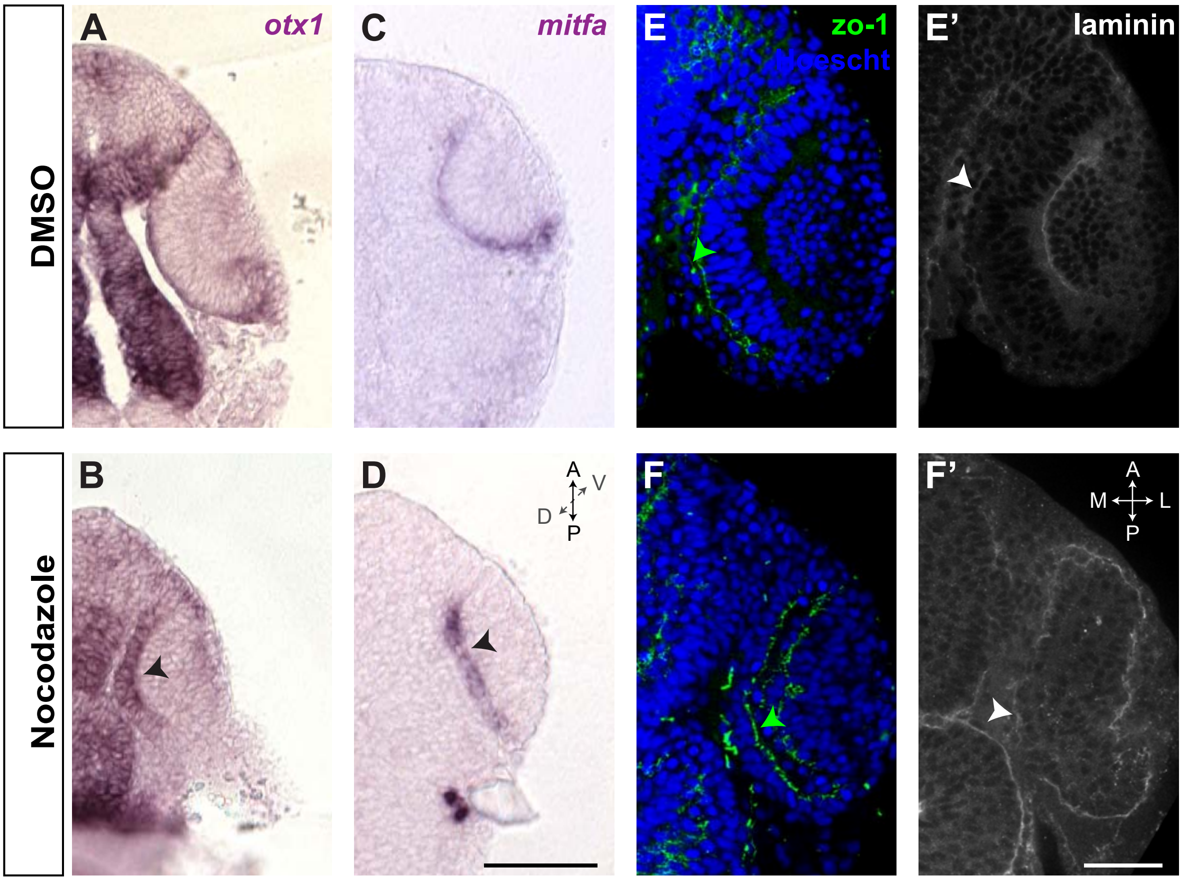

### Fig S4

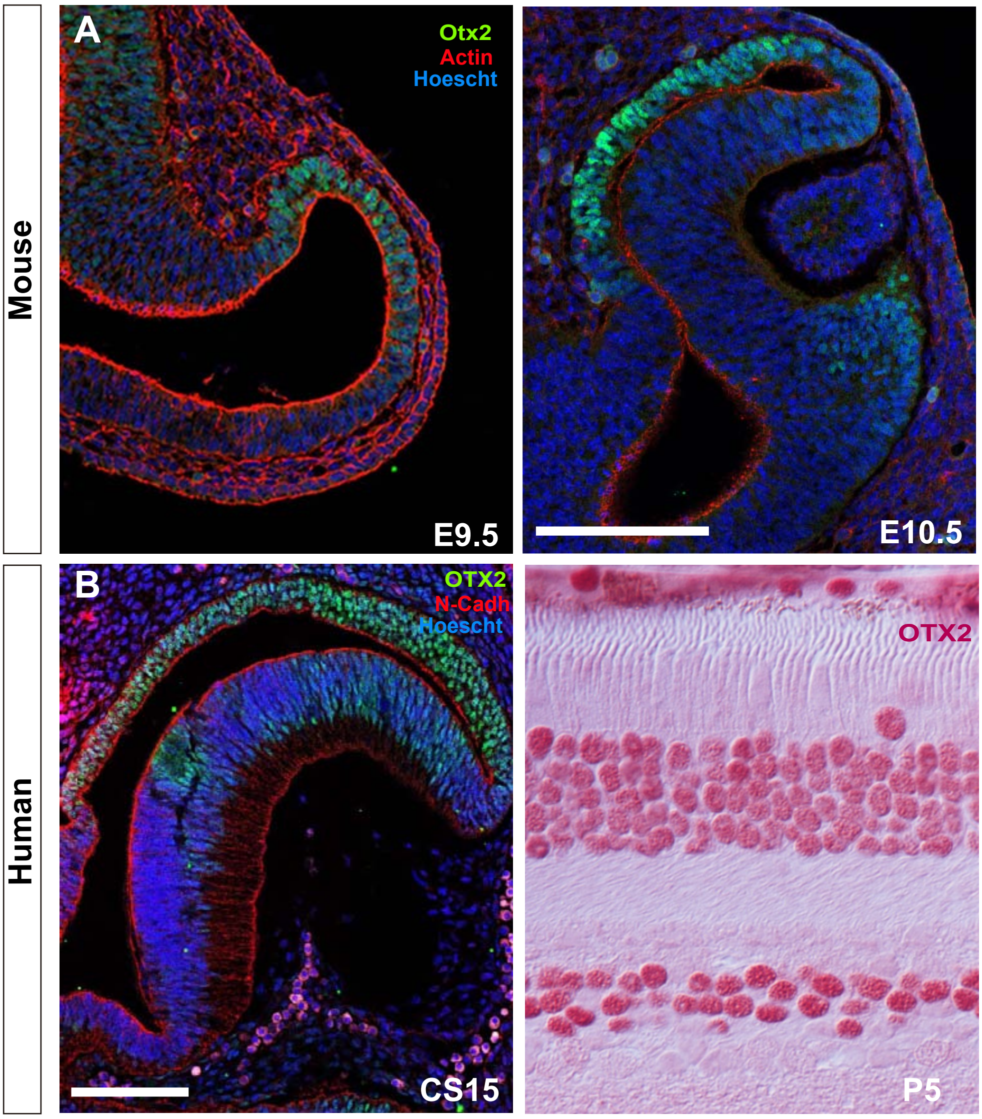
