## Supplementary material for "Stretching of the retinal pigment epithelium contributes to zebrafish optic cup morphogenesis": Table S1

**Table S1. List of primers used in this study**

| **Amplicon** | **Forward** 5’ to 3’ | **Reverse** 5’ to 3’ |
| --- | --- | --- |
| *bhlhe40* Promotor | G AATAGGCTGTCCATGTGGTC | CAAGCCTCAGAAGTAGGACG |
| *bhlhe40* E1 | GTGTAAGGGATGGTCAACAGTG | CAGTTGGGTCAGTTTGAGTTCG |
| *bhlhe40* E2 | GCTTGATGTGTGGACGTTAC | TGTCGCATCACCAGGCTATC |
| *bhlhe40* E3 | GTCCTTGCATGTCAGTGTTTAG | GTAAATCAGCGTTCATCCCAC |
| *bhlhe40*E4 | ACACTGTACGCTTATGGGAG | CCAGAACACCAGGGATAGAC |
| *STMN1* | GGAA**AGGCCT**ATGGCTTCTTCTGATATCCAGGTG | GGAA**AGGCCT**TTAGTCAGCTTCAGTCTCGTC |
| *ccnd1* | GGAA**AGGCCT**GAGGCAGCAAAAAGCATCCAC | GGAA**AGGCCT**CTCGGTCATCAAAGCCACAG |
| *bhlhe40* probe | TGCTACGTAAAAGAAAGCGGG | *TCCATTAACCCTCACTAAAGGGAA*TTCGGGAGCTTATTCAGCAGG |

StuI restriction site introduced for cloning purposes is highlighted in bold.

The sequence of the T3 promoter for probe synthesis is highlighted in italic
